# The *Plasmodium falciparum* Raf Kinase Inhibitor Protein is Essential for Red Blood Cell Invasion and Can Be Functionally Substituted by Host RKIP

**DOI:** 10.64898/2026.04.29.721533

**Authors:** Manish Sharma, Abhisheka Bansal

## Abstract

Raf kinase inhibitor protein (RKIP) plays a key role in regulating critical signaling pathways in higher eukaryotes. We previously demonstrated that *Plasmodium falciparum* RKIP (PfRKIP) modulates the activity of PfCDPK1, a key regulator of red blood cell invasion. Here, we have used a pharmacological approach to investigate the function of PfRKIP. Locostatin, a mammalian RKIP inhibitor, shows a dose-dependent decrease in RBC invasion. Mechanistically, locostatin increases the interaction between PfRKIP and PfCDPK1, thereby sequestering PfCDPK1 in a complex preventing substrate phosphorylation. Surprisingly, PfRKIP could be knocked out from the parasite without any perceptible growth defect. Interestingly, the PfRKIP-deficient parasites show an increase in the import of host RKIP. Like the parasite RKIP, the host RKIP interacts with PfCDPK1 and is associated with high-molecular-weight complexes comprising PfCDPK1 suggesting functional complementation by the host RKIP in the PfRKIP null parasites. As expected, locostatin shows similar inhibitory effect on the knock-out parasites as the WT. Our study reveals a novel host-parasite interaction wherein the parasite co-opts host proteins to maintain critical signaling pathways. Targeting the PfRKIP signaling axis, along with the host proteins, represents a potential strategy for anti-malarial drug development that is conceptually less prone to parasite-driven resistance development.

## INTRODUCTION

Malaria continues to pose a severe burden on global health, particularly in developing countries^1^. The emergence of *Plasmodium falciparum* strains exhibiting delayed parasite clearance and resistance to artemisinin-based combination therapies underscores the urgent need to identify novel molecular targets and develop improved strategies for malaria control^2–4^. A detailed understanding of the signaling pathways that regulate red blood cell (RBC) invasion is critical for devising such interventions.

Raf Kinase Inhibitor Protein (RKIP) is a well-characterized regulator of the MEK/ERK signaling cascade in higher eukaryotes^5,6^. Under basal conditions, RKIP sequesters Raf1, preventing its interaction with MEK and thereby inhibiting downstream ERK activation. Pharmacological disruption of this interaction using locostatin, a small-molecule RKIP inhibitor, releases Raf1 from RKIP, resulting in activation of the MEK/ERK pathway^5,7,8^. Interestingly, canonical components of the MEK/ERK pathway are absent in *P. falciparum*, suggesting that PfRKIP has diverged to perform unique functions distinct from its mammalian ortholog.

PfRKIP contains a phosphatidylethanolamine-binding (PEBP) domain and interacts with various physiologically relevant lipids, implicating it in membrane-associated signaling^9^. We have previously demonstrated that PfRKIP directly interacts with PfCDPK1, a calcium-dependent protein kinase essential for RBC invasion, within mature schizonts, and modulates its kinase activity^10^. Notably, phosphorylation of PfRKIP by PfCDPK1 in the presence of substrate reduces their interaction, thereby facilitating the transphosphorylation of the substrate by PfCDPK1.

PfCDPK1 is a canonical member of the calcium-dependent protein kinase family, which orchestrates multiple processes during parasite development^11–14^. Its substrates include proteins of the inner membrane complex and the glideosome, both of which are essential components of the actin-myosin motor required for parasite motility and RBC invasion^15,16^. Furthermore, PfCDPK1-mediated phosphorylation of regulatory proteins, such as PKAr, is critical for invasion, and the kinase also facilitates microneme discharge, a pivotal step in the sequential events that drive RBC entry^12,13^. Despite extensive knowledge of PfCDPK1 function and downstream substrates, the molecular mechanisms governing its regulation remain incompletely understood.

Host RKIP (HsRKIP), also referred to as phosphatidylethanolamine-binding protein 1 (PEBP1), exhibits a lipid interaction profile distinct from PfRKIP under *in vitro* conditions ^10^. In mammalian cells, HsRKIP interacts with numerous cytoplasmic proteins, modulating key signaling pathways involved in proliferation, migration, and apoptosis^5,17,18^. RKIP negatively regulates the NFκB pathway by inhibiting upstream kinases^17^, and perturbations in RKIP expression or function are associated with various human pathologies^19–21^. RKIP exists in multiple functional states, determined in part by phosphorylation, which modulates its capacity to interact with diverse protein partners^22,23^.

During intraerythrocytic development, *P. falciparum* imports multiple host proteins and nutrients from the RBC cytoplasm to meet its metabolic demands^24,25^. Hemoglobin uptake, a major nutrient source, generates significant oxidative stress and lipid peroxidation. To counteract these effects, the parasite imports lipid repair enzymes, including peroxiredoxin 6 (PRDX6)^26^. Similarly, HsRKIP, an abundant RBC protein, is internalized by the parasite during its growth^26–28^. Despite consistent observations of HsRKIP import, the functional significance of this process remained largely unexplored. It is interesting to note that HsRKIP also interacts with PfCDPK1 albeit with a lower affinity^10^.

In this study, we demonstrate that treatment of mature schizonts with locostatin leads to a dose-dependent inhibition of RBC invasion. Mechanistically, locostatin increases the interaction between PfRKIP and PfCDPK1, in contrast to its inhibitory effect on the HsRKIP-Raf1 interaction in mammalian cells. The sequestration of PfCDPK1 within this complex reduces its availability for downstream substrate phosphorylation, providing a plausible explanation for the impaired invasion. Surprisingly, complete disruption of *pfrkip* does not affect asexual proliferation. Intriguingly, the *pfrkip* knock-out (*Δpfrkip*) parasites exhibit enhanced import of HsRKIP during late developmental stages. We further show that HsRKIP engages PfCDPK1 in late-stage parasites and forms high-molecular-weight complexes reminiscent of those observed with PfRKIP. Collectively, these findings underscore the divergent evolution of PfRKIP as a critical regulator of PfCDPK1-mediated signaling, distinct from canonical MEK/ERK regulation in higher eukaryotes.

## RESULTS

### Pharmacological inhibition of PfRKIP blocks RBC invasion

We previously demonstrated that PfRKIP and PfCDPK1 co-localize at the apical end of free and developing merozoites within mature schizonts, and that PfRKIP interacts with and regulates the activity of PfCDPK1^10^. To investigate the functional implications of PfRKIP inhibition on asexual proliferation of the parasite, we employed locostatin, a small-molecule inhibitor of mammalian RKIP known to disrupt RKIP-Raf1 interactions, thereby activating the MAPK signaling cascade^5,7,8^. Given the established role of PfCDPK1 in RBC invasion, we hypothesized that locostatin-mediated modulation of PfRKIP could impair the parasite’s ability to invade RBCs. To test this, synchronized mature schizonts (38–42 hours post-invasion, HPI) were exposed to increasing concentrations of locostatin, and the formation of ring-stage parasites in the subsequent cycle was assessed microscopically by Giemsa-stained thin smears. Treatment with locostatin resulted in a dose-dependent reduction in the number of newly formed ring-stage parasites (Fig. 1a; n=3; p<0.05, RM one-way ANOVA, Dunnett’s multiple comparisons test). These findings were corroborated using an independent, quantitative assay; an automated SYBR Green I–based analysis showed a similar concentration-dependent decrease in parasite-derived fluorescence, consistent with reduction in the parasite proliferation (Fig. S1).

**Figure 1.**
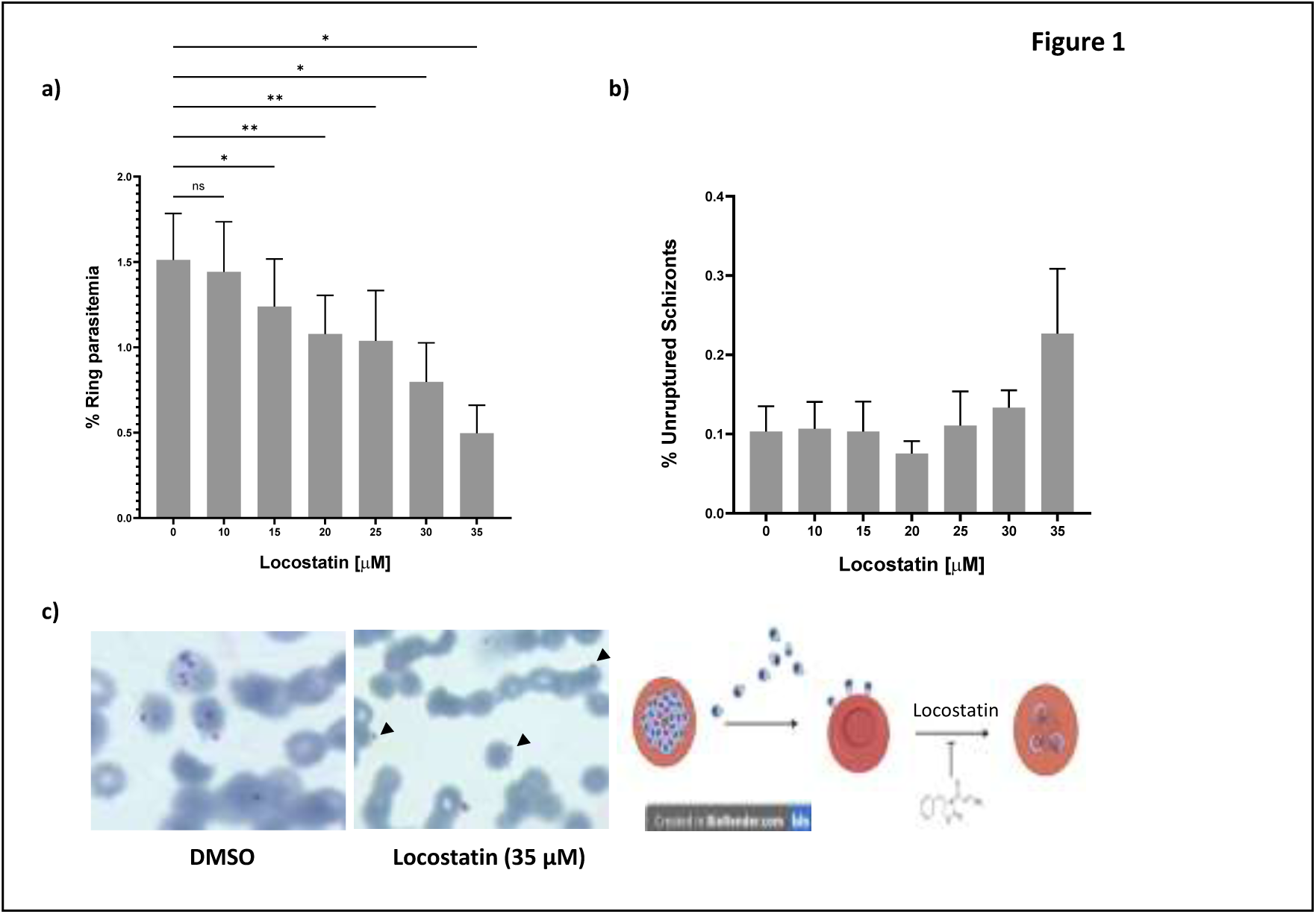
Pharmacological inhibition of PfRKIP blocks the invasion of red blood cells by the malaria parasite. **a)** To assess the effect of PfRKIP inhibition on the invasion of red blood cells (RBCs), we used a specific inhibitor of mammalian RKIP called locostatin. Highly synchronized schizonts of WT parasites were treated with varying concentrations of locostatin. The ring-stage parasites were counted 12–14 h post treatment through Giemsa stained smears. The number of ring-stage parasites decreased in a dose-dependent manner with increasing concentration of locostatin. The percent (%) ring parasitemia is plotted on the Y-axis against locostatin concentration (in μM) on the X-axis. The data is obtained from 3 independent biological experiments counted in duplicate (n=3; *p<0.05; **p<0.005; ns-not significant; RM one-way ANOVA). Error bars represent the standard error of the mean. **b)** Locostatin does not show significant defect on the egress of merozoites. The unruptured schizonts were counted in all the locostatin treatment conditions along with the rings. The % unruptured schizonts on the Y-axis are plotted against different concentrations of locostatin on the X-axis. The difference in the % unruptured schizonts is not statistically significant (n=3 independent biological experiments, p>0.05, RM one-way ANOVA). The error bars represent the standard error of the mean. **c)** Representative images of Giemsa smears (100X magnification) show defect in the parasite invasion of RBCs. In the locostatin (35 μM) condition, less ring infected parasites are seen than the control (DMSO). Merozoites attached to the RBCs are seen in the locostatin condition (black arrowhead) suggesting defect in the invasion process. (lower right) Scheme showing the inhibitory effect of locostatin on the parasite invasion of RBC. Egress of merozoites from an infected RBC is not affected by locostatin whereas the subsequent invasion of a fresh RBC is blocked.

To distinguish whether this defect reflected impaired egress or invasion, we quantified unruptured schizonts under each treatment condition. Up to 30 μM locostatin, the proportion of unruptured schizonts remained comparable to untreated controls (Fig. 1b). Above 30 μM locostatin, a non-significant trend toward increased unruptured schizonts was observed (Fig. 1b), indicating that locostatin predominantly affects the invasion process rather than schizont rupture. Importantly, free merozoites attached to the RBC surface were observed in the Giemsa smears under locostatin treatment condition compared to rings in the control (Fig. 1c), further suggesting that locostatin inhibits the RBC invasion by merozoites rather than the egress process. Collectively, these data indicate that locostatin selectively impairs RBC invasion by *P. falciparum*, likely through modulation of PfRKIP-PfCDPK1 interaction, without a significant effect on egress.

### Locostatin enhances the interaction of PfRKIP with PfCDPK1

We sought to elucidate the mechanism underlying locostatin-mediated inhibition of RBC invasion. Locostatin inhibits HsRKIP by alkylating His96^29^, an event that induces conformational changes in RKIP and alters its interactions with partner proteins, thereby modulating downstream signaling pathways. We previously demonstrated that PfRKIP interacts with PfCDPK1 both *in vitro* and within the parasite^10^. To determine whether locostatin influences the interaction between PfRKIP and PfCDPK1, we employed an ELISA-based binding assay.

PfRKIP pre-incubated with increasing concentrations of locostatin was allowed to interact with PfCDPK1 immobilized on a 96-well plate. Bound PfRKIP was detected using anti-PfRKIP antibody, and absorbance was measured at 492 nm (OD_492 nm_). Contrary to our expectation, locostatin-treated PfRKIP exhibited significantly enhanced binding with PfCDPK1 compared to untreated PfRKIP across all concentrations tested (Fig. 2a; n = 3; p<0.05, multiple paired t-test). Notably, when PfRKIP was not pre-incubated with locostatin but instead added simultaneously with other assay components, no change in its interaction with PfCDPK1 was observed (Fig. S2a). These results indicate that pre-incubation of PfRKIP with locostatin is essential for enhanced binding with PfCDPK1, likely due to alkylation of a specific histidine residue in PfRKIP, analogous to the modification of His96 in HsRKIP^29,30^. Together, these data suggest that locostatin may inhibit RBC invasion by enhancing the interaction between PfRKIP and PfCDPK1.

**Figure 2.**
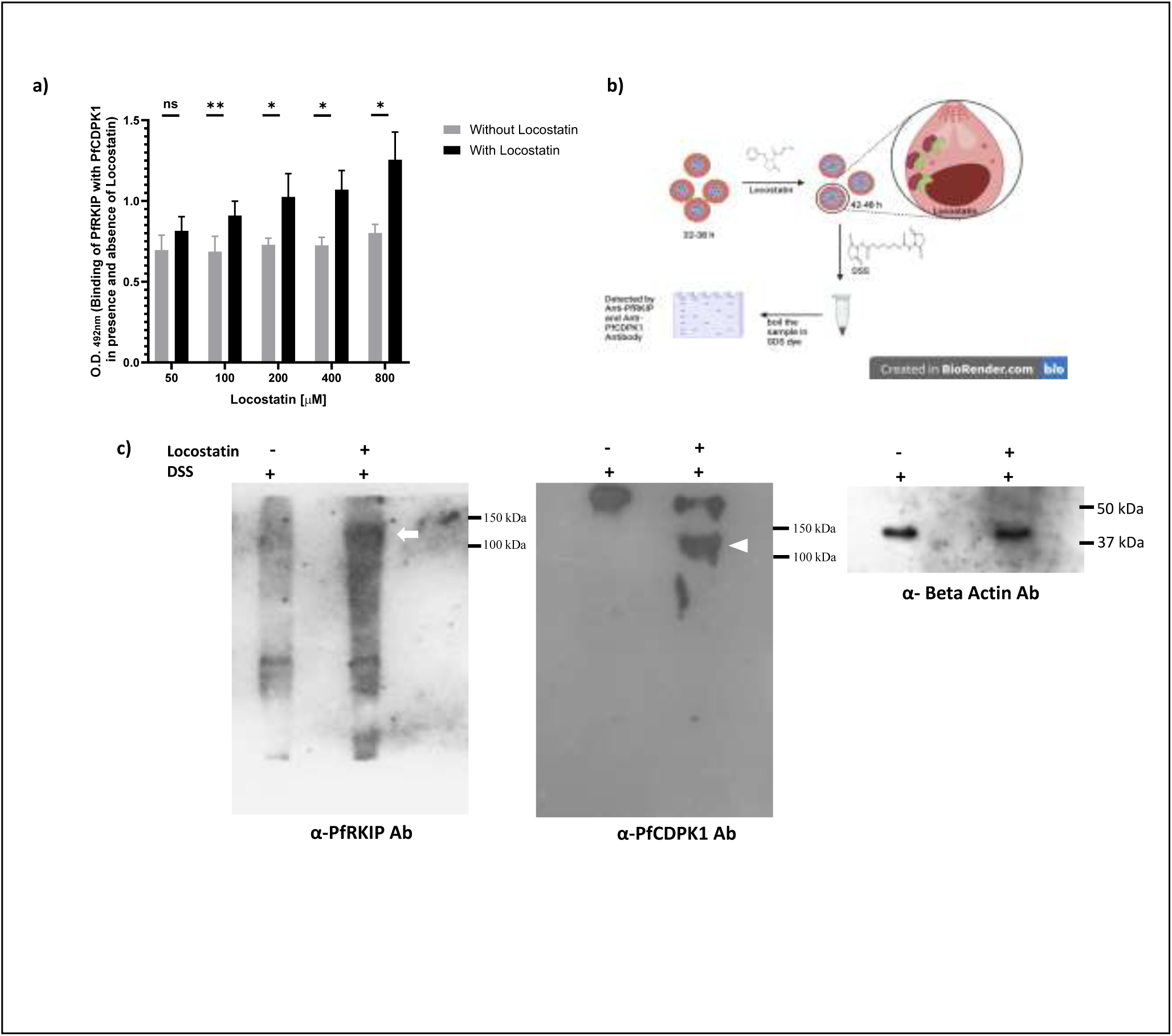
Locostatin increases the interaction of PfRKIP with PfCDPK1. **a)** Locostatin mediated modification of recombinant PfRKIP increases its interaction with recombinant PfCDPK1. Purified PfRKIP was treated with different concentrations of locostatin and allowed to interact with PfCDPK1 immobilized on a 96-well plate. Locostatin-treated PfRKIP show higher binding with PfCDPK1 compared to without locostatin condition at all the concentrations of the compound. The Optical Density (OD_492 nm_) representing the binding of PfRKIP with PfCDPK1 in the presence and absence of locostatin on the Y-axis is plotted against the concentrations of locostatin (in μM) on the X-axis. The data is obtained from three independent biological experiments performed in duplicate (n=3, *p<0.05, **p<0.005, ns-non-significant; multiple paired t-test). Error bars represent the standard error of the mean. **b & c)** Treatment of the parasites with locostatin increases the interaction of PfRKIP and PfCDPK1. **b)** Schematic representation of the experimental setup to test the effect of locostatin on the interaction of PfRKIP and PfCDPK1 within the parasite. Late-stage trophozoite (32-38 HPI) were treated with or without locostatin and allowed to progress to 42-48 HPI. The parasites were treated with DSS (disuccinimidyl suberate), a cell permeable, irreversible crosslinking reagent that covalently links all the interacting proteins. The DSS treated samples were processed further followed by Western blot with anti-PfRKIP and anti-PfCDPK1 antibodies. **c)** Parasites treated with locostatin show increase in the intensity of a high molecular weight complex containing PfRKIP and PfCDPK1. Lysates prepared from the parasites treated with or without locostatin followed by DSS cross-linking were separated on SDS-PAGE followed by Western blot using anti-PfRKIP and anti-PfCDPK1 antibodies. Both the antibodies show the detection of a high molecular weight complex between ∼100 kDa and ∼150 kDa (white arrow). The intensity of the high molecular weight complex is more in the locostatin-treated samples than the untreated control, indicating enhanced interaction between PfRKIP and PfCDPK1 in the presence of locostatin. Western with anti-β-actin antibody show equal protein loading in both the samples.

To validate these findings in a cellular context, we examined the effect of locostatin on the PfRKIP–PfCDPK1 interaction within parasites. We used a PfRKIP-overexpressing parasite line (RKIP-cMyc)^10^. Late-stage parasites (32–38 HPI) were treated with locostatin for 10 h, followed by covalent cross-linking of protein–protein interactions using disuccinimidyl suberate (DSS), a cell-permeable, irreversible crosslinker (Fig. 2b). Protein extracts from locostatin-treated and untreated parasites were analyzed by Western blotting using anti-PfRKIP and anti-PfCDPK1 antibodies.

In locostatin-treated parasites, probing with anti-PfRKIP antibody revealed a high-molecular-weight complex between ∼100 and 150 kDa (Fig. 2c, white arrow). Following stripping and re-probing with anti-PfCDPK1 antibody, the same complex was detected, confirming the presence of PfCDPK1 within this assembly (Fig. 2c). Importantly, this complex was not detected in the without locostatin condition. In addition, a larger PfCDPK1-containing complex (>150 kDa) was observed exclusively with anti-PfCDPK1 antibody. Notably, the intensity of the >150 kDa complex was reduced in locostatin-treated parasites compared to the untreated control, suggesting that locostatin promotes the sequestration of PfCDPK1 into a PfRKIP-containing complex, thereby limiting its availability to participate in other higher-order protein assemblies. In contrast, in the absence of locostatin, PfCDPK1 appears to form higher-molecular-weight complexes with proteins other than PfRKIP. A parallel blot probed with anti-β-actin antibody confirmed equal protein loading across samples (Fig. 2c).

We previously reported that ATP reduces the interaction between PfRKIP and PfCDPK1^10^. To assess whether phosphorylation similarly affects the interaction of locostatin-treated PfRKIP with PfCDPK1, recombinant PfRKIP at varying concentrations was pre-treated with 800 μM locostatin and incubated with PfCDPK1 in the presence or absence of ATP. Bound PfRKIP was detected using anti-PfRKIP antibody. Consistent with our prior observations, locostatin-treated PfRKIP exhibited significantly reduced interaction with PfCDPK1 in the presence of ATP compared to ATP-free conditions (Fig. S2b, n = 3, p<0.05, paired t-test), indicating that phosphorylation diminishes the PfRKIP–PfCDPK1 interaction irrespective of locostatin treatment^10^ (Fig. S2b).

Collectively, our findings support a model in which locostatin enhances the interaction between PfRKIP and PfCDPK1, sequestering PfCDPK1 into a specific complex that prevents the formation of higher-order assemblies potentially required for successful RBC invasion.

### Locostatin-modified RKIP sequesters PfCDPK1 and reduces substrate phosphorylation

Given that locostatin-treated PfRKIP exhibits enhanced interaction with PfCDPK1, we next examined whether sequestration of PfCDPK1 by PfRKIP affects the transphosphorylation activity of PfCDPK1. To address this, we performed an *in vitro* kinase assay using recombinant PfCDPK1 and myelin basic protein (MBP), used as an exogenous substrate. PfRKIP pre-treated with either locostatin (modified PfRKIP) or DMSO (unmodified PfRKIP) was included in the kinase reaction mixture.

Following the kinase reaction, the samples were resolved by SDS–PAGE, and phosphorylated proteins were visualized using Pro-Q Diamond phosphoprotein stain. In the presence of modified PfRKIP, phosphorylation of MBP was markedly reduced compared to the control condition containing unmodified PfRKIP (Fig. 3a, white arrow). The Pro-Q Diamond–stained gel was subsequently counterstained with Coomassie Brilliant Blue R-250 to assess total protein loading, which was comparable between the two conditions (Fig. 3a, right). Quantification of phosphorylated MBP was performed using ImageJ, and Pro-Q Diamond signal intensity was normalized to the corresponding Coomassie-stained MBP band. The reduction in MBP transphosphorylation observed in the presence of modified PfRKIP was statistically significant (Fig. 3b; n=3, p = 0.0086, unpaired t-test).

**Figure 3.**
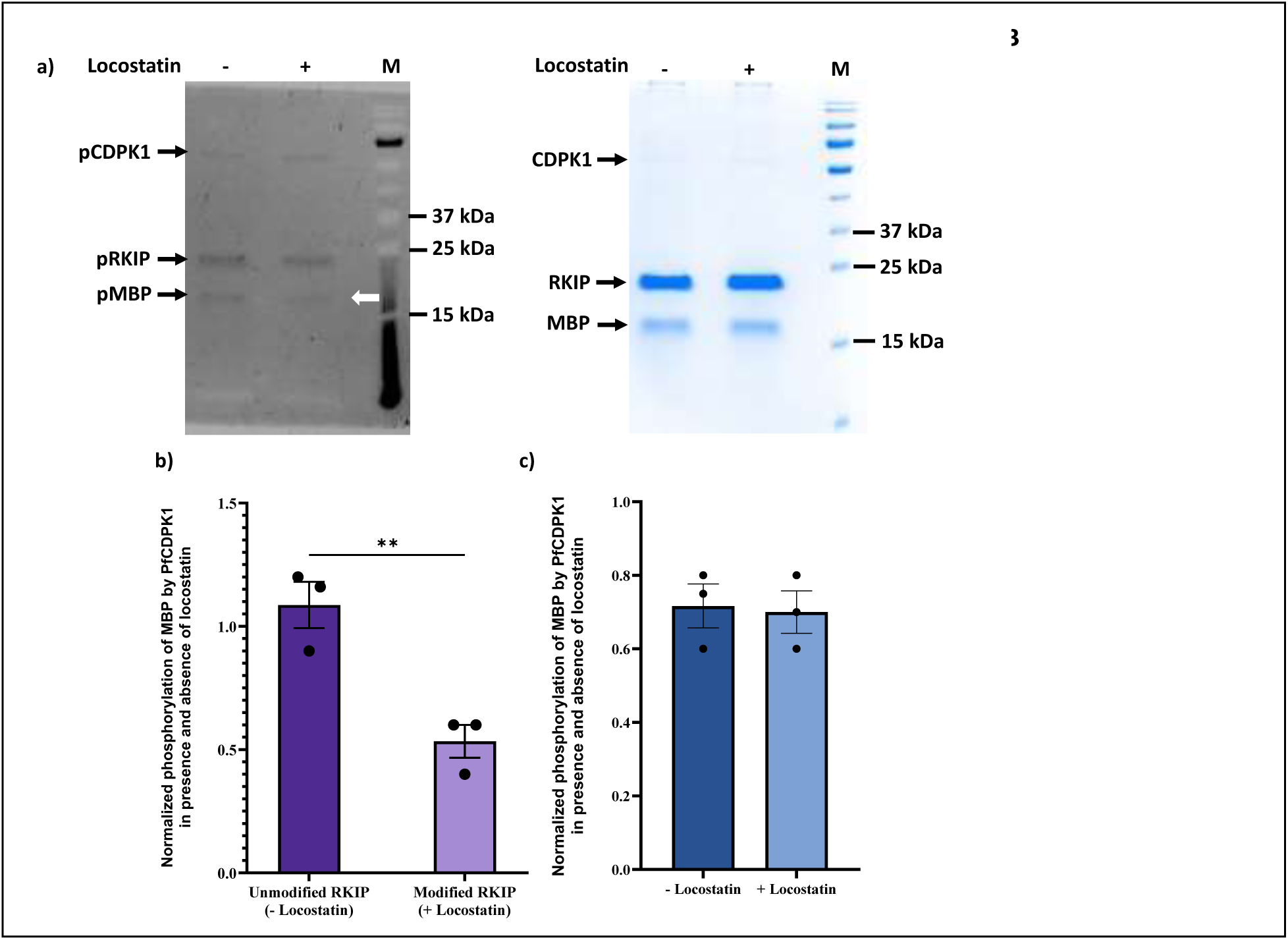
Locostatin-modified PfRKIP decreases the trans-phosphorylation activity of PfCDPK1. **a)** PfRKIP, pre-incubated with either locostatin or DMSO was used to set up an *in vitro* kinase assay with PfCDPK1 along with an exogenous substrate, Myelin Basic Protein (MBP). Substrate-level phosphorylation of MBP by PfCDPK1 was analyzed by resolving the reaction mixture on SDS-PAGE followed by staining with Pro-Q Diamond phosphoprotein stain. The Pro-Q Diamond-stained gel shows a reduction in the phosphorylation of exogenous MBP (indicated by a white arrow) in the locostatin-treated condition compared to the untreated (DMSO) control. (Right) The same gel was subsequently stained with Coomassie Brilliant Blue R250. M- unstained protein ladder. **b)** The intensity of the phosphorylated MBP band was quantified using ImageJ software and normalized with the Coomassie stained MBP protein. The histogram shows the normalized phosphorylation of MBP by PfCDPK1 on the Y-axis in the presence of unmodified (- Locostatin) and modified (+ Locostatin) PfRKIP on the X-axis. In the presence of locostatin, the phosphorylation of MBP is significantly reduced (n=3, **p=0.0086, unpaired t-test). The error bars represent the standard error of the mean. **c)** Locostatin *per se* has no effect on the transphosphorylation activity of PfCDPK1. *In vitro* kinase assay with PfCDPK1 was set up in the presence and absence of locostatin using MBP as an exogenous substrate. Recombinant PfRKIP was not included in the reaction mixture. The histogram represents the normalized phosphorylation of MBP by PfCDPK1 on the Y-axis in the presence and absence of locostatin on the X-axis. Locostatin does not affect the kinase activity of PfCDPK1 (n=3, p>0.05; unpaired t-test). The error bars represent the standard error of the mean.

To determine whether locostatin directly affects the kinase activity of PfCDPK1, we conducted *in vitro* kinase assays in the absence of PfRKIP. Under these conditions, the transphosphorylation of MBP by PfCDPK1 was not diminished in the presence of locostatin compared to control reactions lacking locostatin (Fig. 3c and Fig. S2c). Consistently, normalized MBP phosphorylation levels were similar between locostatin-treated and untreated reactions (Fig. 3c). These findings indicate that locostatin *per se* has no inhibitory effect on PfCDPK1 kinase activity. Taken together, these results suggest that the enhanced interaction between locostatin-modified PfRKIP and PfCDPK1 likely reduces the pool of free PfCDPK1 available for substrate phosphorylation.

### PfRKIP is dispensable for asexual replication of the parasite under *in vitro* conditions

A genome-wide mutagenesis study previously reported that *pfrkip* is non-mutable within its coding sequence, suggesting that the gene may be essential for parasite growth within RBCs^31^. To directly test the indispensability of *pfrkip* for parasite growth, we disrupted the endogenous *pfrkip* locus using CRISPR/Cas9-mediated gene editing.

A single plasmid, *pL6-Cas9-pfrkipko*, containing a guide RNA targeting the PEBP domain of PfRKIP, the Cas9 endonuclease, and a homologous repair template, was constructed and used for parasite transfection (Fig. 4a). Following WR99210 selection, viable parasites were detected approximately three weeks post-transfection. Clonal parasites were subsequently obtained by limiting dilution and analyzed for targeted disruption of the *pfrkip* locus by PCR using locus-specific primers. Amplification of the full-length locus using F3/R1 primer pair yielded amplicons of 1305 bp and 2994 bp in wild-type (WT) and *Δpfrkip* parasites, respectively (Fig. 4b). The increased amplicon size in the *Δpfrkip* parasites corresponds to replacement of the PEBP domain with the human dihydrofolate reductase (hDHFR) selection cassette.

**Figure 4.**
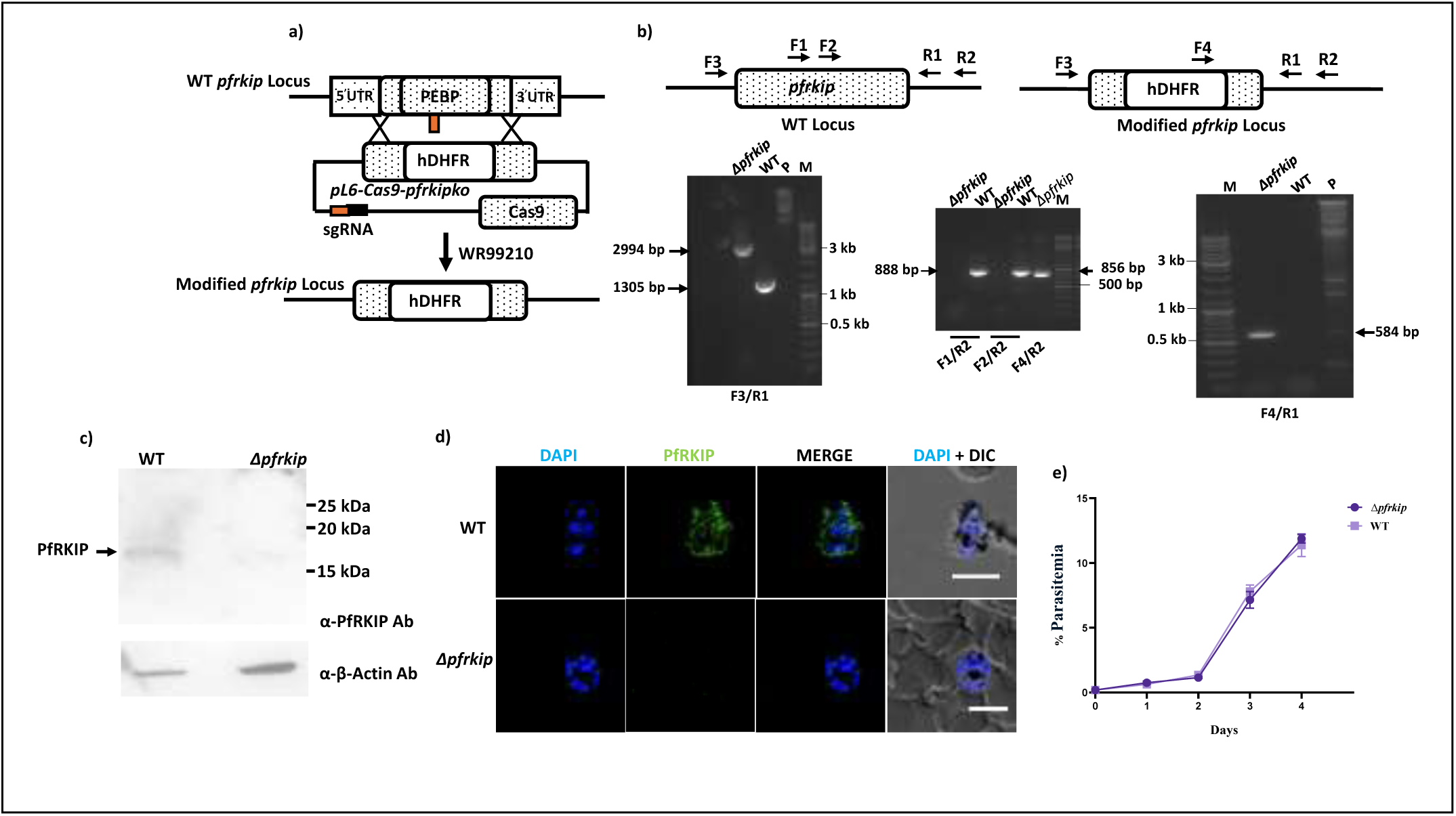
PfRKIP is dispensable for the asexual replication of the parasite. **a)** Scheme representing the strategy for the disruption of *pfrkip* gene. The PEBP domain of PfRKIP is replaced with the human dihydrofolate reductase (hDHFR) cassette through CRISPR/Cas9 gene editing. The scheme shows the guide sequence as a solid red box along with the tracrRNA (black box). The single guide RNA (sgRNA) consists of the guide and the tracer RNA. The *pL6-Cas9-pfrkipko* plasmid contains all the essential elements required for the desired editing: sgRNA, hDHFR resistance cassette, Cas9, and the homology region. After the desired recombination, the *pfrkip* locus is disrupted by the substitution of the PEBP domain with the hDHFR expression cassette. **b)** PCR verification of *pfrkip* disruption. PCR with specific primer pairs show successful knock-out of *pfrkip* gene. Oligos, F1 and F2 corresponding to the PEBP DNA sequence show specific amplification only in the WT parasites when combined with the R2 oligo. No amplification is seen in the *pfrkip* KO (Δ*pfrkip*) parasite. The F4/R2 primer pair produces an amplicon of 848 bp in the Δ*pfrkip* parasites confirming the integrity of the template DNA used for the PCR reaction. The F3/R1 oligo pair produces amplicons of 1305 and 2994 bp in the WT and the Δ*pfrkip* parasites, respectively. The *pL6-Cas9-pfrkipko* (P) plasmid is taken as a negative control. The F4/R1 primer pair produces an amplicon of 584 bp only in the Δ*pfrkip* parasite as F4 is complementary to the hDHFR sequence while R1 is complementary to the 3’ UTR region of *pfrkip* gene. M- GeneRuler DNA Ladder Mix. **c)** PfRKIP protein is not present in the Δ*pfrkip* parasites. Parasite lysates prepared from the mature schizonts of Δ*pfrkip* and WT parasites were analyzed through Western blot using anti-PfRKIP antibody. In the WT parasites, a band of the expected size (∼ 21.8 kDa), corresponding to PfRKIP, is seen, that is absent in the Δ*pfrkip* parasite lysate. A parallel blot was probed with the anti-Beta actin antibody as a loading control. **d)** Immunofluorescence assay (IFA) confirms the absence of PfRKIP in the Δ*pfrkip* parasite. The WT and the Δ*pfrkip* parasites were probed with the anti-PfRKIP antibody followed by an appropriate secondary antibody conjugated to Alexafluor 488. In the WT parasites, expected fluorescence of PfRKIP is observed. No PfRKIP specific fluorescence is detected in the Δ*pfrkip* parasite, further confirming the absence of RfRKIP in the Δ*pfrkip* parasites. The parasites were counterstained with DAPI, a nuclear stain. White error bars in the panel, DAPI + DIC (differential interference contrast) correspond to 5 μm. **e)** The *pfrkip* knock-out parasites grow similarly to the WT parasites. Asexual growth of highly synchronized WT and Δ*pfrkip* parasites were compared for two consecutive cycles by counting the Giemsa smears every 24 h for the period of 4 days. The growth curve of Δ*pfrkip* parasites closely mirrors the WT parasites, indicating that PfRKIP is not essential for the parasite proliferation under *in vitro* conditions. The graph represents the % parasitemia on the Y-axis for the corresponding days on the X-axis. The error bars represent the standard error of the mean (n=2 independent biological experiments). The graph is plotted using GraphPad Prism version 10.6.1.

To further validate the disruption of *pfrkip*, diagnostic PCRs were performed using additional primer combinations. Primer pairs F1/R2 and F2/R2, spanning the deleted PEBP domain and the downstream 3′ untranslated region (UTR), produced amplicons of the expected sizes (888 bp and 856 bp, respectively) in WT parasites but not in the *Δpfrkip* parasites (Fig. 4b). Integrity of the genomic template in the *Δpfrkip* parasites was confirmed using the F4/R2 primer pair, which yielded an amplicon of the expected size (848 bp) (Fig. 4b). Integration of the hDHFR cassette at the *pfrkip* locus was further confirmed using the F4/R1 primer pair, which generated a 584 bp amplicon exclusively in *Δpfrkip* parasites and not in the WT parasites (Fig. 4b). Collectively, these PCR analyses confirm successful disruption of *pfrkip* via replacement of the PEBP domain with the hDHFR cassette.

To confirm loss of *pfrkip* expression at the transcript level, semi-quantitative RT-PCR was performed using primers corresponding to the deleted region of *pfrkip*. No amplification was observed in the *Δpfrkip* parasites, whereas products of the expected sizes of 325 bp and 590 bp were detected in WT parasites with RKIPS96A/PfRKIPG4T1_R and RKIPV5F1/ PfRKIPG4T1_R primer pairs, respectively (Fig. S3a). Amplification of a remnant region of *pfrkip* using the RKIPFRT/RKIPRRT primer pair yielded an amplicon of the expected size (118 bp) in *Δpfrkip* parasites, confirming cDNA integrity (Fig. S3a). Complete disruption of PfRKIP was further validated at the protein level by Western blot analysis of mature schizonts, the stage at which PfRKIP is maximally expressed^10,32^. A PfRKIP-specific band was detected in WT parasites but was absent in *Δpfrkip* parasites (Fig. 4c). Reprobing the blot with anti-β-actin antibodies confirmed equal protein loading (Fig. 4c). Consistently, immunofluorescence assays performed on schizont-stage parasites showed PfRKIP-specific staining in WT parasites, whereas no signal was detected in *Δpfrkip* parasites (Fig. 4d). Together, these results conclusively demonstrate successful knockout of *pfrkip*.

Following confirmation of *pfrkip* disruption, we assessed whether loss of PfRKIP impacts parasite asexual growth. WT and *Δpfrkip* parasites were synchronized with two consecutive sorbitol treatments and cultured for four days. Parasite growth was monitored daily by Giemsa-stained thin blood smears. *Δpfrkip* parasites exhibited growth kinetics comparable to WT parasites (Fig. 4e). These findings were independently validated using a SYBR Green I–based fluorescence assay, which likewise showed no difference in growth between WT and *Δpfrkip* parasites (Fig. S3b). Collectively, these data indicate that PfRKIP is dispensable for asexual intraerythrocytic growth under *in vitro* culture conditions.

*P. falciparum* encodes another PEBP domain–containing protein, PF3D7_0303900, which is also maximally expressed during the schizont stage (PlasmoDB). To determine whether *Δpfrkip* parasites compensate for the loss of PfRKIP by upregulating PF3D7_0303900, we examined its transcript level in schizont-stage parasites using gene specific primer pair, PEBP3900FP/ PEBP3900RP (612 bp amplicon). Expression of PF3D7_0303900 was comparable between WT and *Δpfrkip* parasites (Fig. S4). Additionally, given the established interaction between PfRKIP and PfCDPK1 and previous observations that disruption of *pfcdpk1* leads to increased *pfrkip* expression^10,11^, we assessed *pfcdpk1* transcript level in *Δpfrkip* parasites. No significant difference in *pfcdpk1* expression was observed between WT and *Δpfrkip* parasites (Fig. S4). These findings suggest that *Δpfrkip* parasites do not compensate for PfRKIP loss by transcriptional upregulation of related PEBP proteins or PfCDPK1, implying either the existence of alternative adaptive mechanisms or that PfRKIP is redundant for parasite growth within RBCs under *in vitro* conditions.

### Δ*pfrkip* parasites show increased import of host RKIP

HsRKIP is abundantly expressed in RBCs and is imported by *P. falciparum* during intraerythrocytic development^26–28^. However, the functional significance of HsRKIP import by the parasite remains unexplored. We previously demonstrated that HsRKIP interacts with PfCDPK1, albeit with more than ∼23-fold lower affinity than PfRKIP^10^. These observations provided a rationale to examine whether loss of PfRKIP influences the import of HsRKIP in the parasite.

To specifically detect HsRKIP, we employed a commercial anti-HsRKIP antibody (c-anti-HsRKIP Ab) that is generated against a unique sequence at the C-terminus. The specificity of this antibody was first validated by Western blot analysis using recombinant PfRKIP and HsRKIP proteins. The c-anti-HsRKIP Ab specifically detected recombinant HsRKIP but did not recognize PfRKIP (Fig. S5a). An RBC lysate included as a positive control showed a band corresponding to HsRKIP at the expected molecular weight (∼21 kDa) (Fig. S5a). Equal loading of recombinant PfRKIP and HsRKIP proteins was confirmed using an anti-His antibody, as both proteins contain a C-terminal 6×His tag^10^ (Fig. S5a). These results confirm the specificity of the c-anti-HsRKIP Ab for HsRKIP with no detectable cross-reactivity toward PfRKIP.

Next, we optimized conditions to generate parasite lysates devoid of host RBC contamination to exclude the possibility of HsRKIP carry-over from the host cytoplasm. Schizont-stage WT parasites treated with or without saponin were analyzed by Western blot using an anti-hemoglobin antibody. Hemoglobin was undetectable in saponin-treated samples, confirming efficient removal of host RBC contents under the conditions employed (Fig. S5b). Notably, a band corresponding to HsRKIP was detected in saponin-treated parasite lysates using the c-anti-HsRKIP Ab (Fig. 5a and Fig. S5b), whereas HsRKIP signal was markedly stronger in saponin-untreated samples, consistent with the presence of intact RBCs (Fig. S5b). Comparable protein loading between samples was confirmed using anti-β-actin antibodies (Fig. S5b).

**Figure 5.**
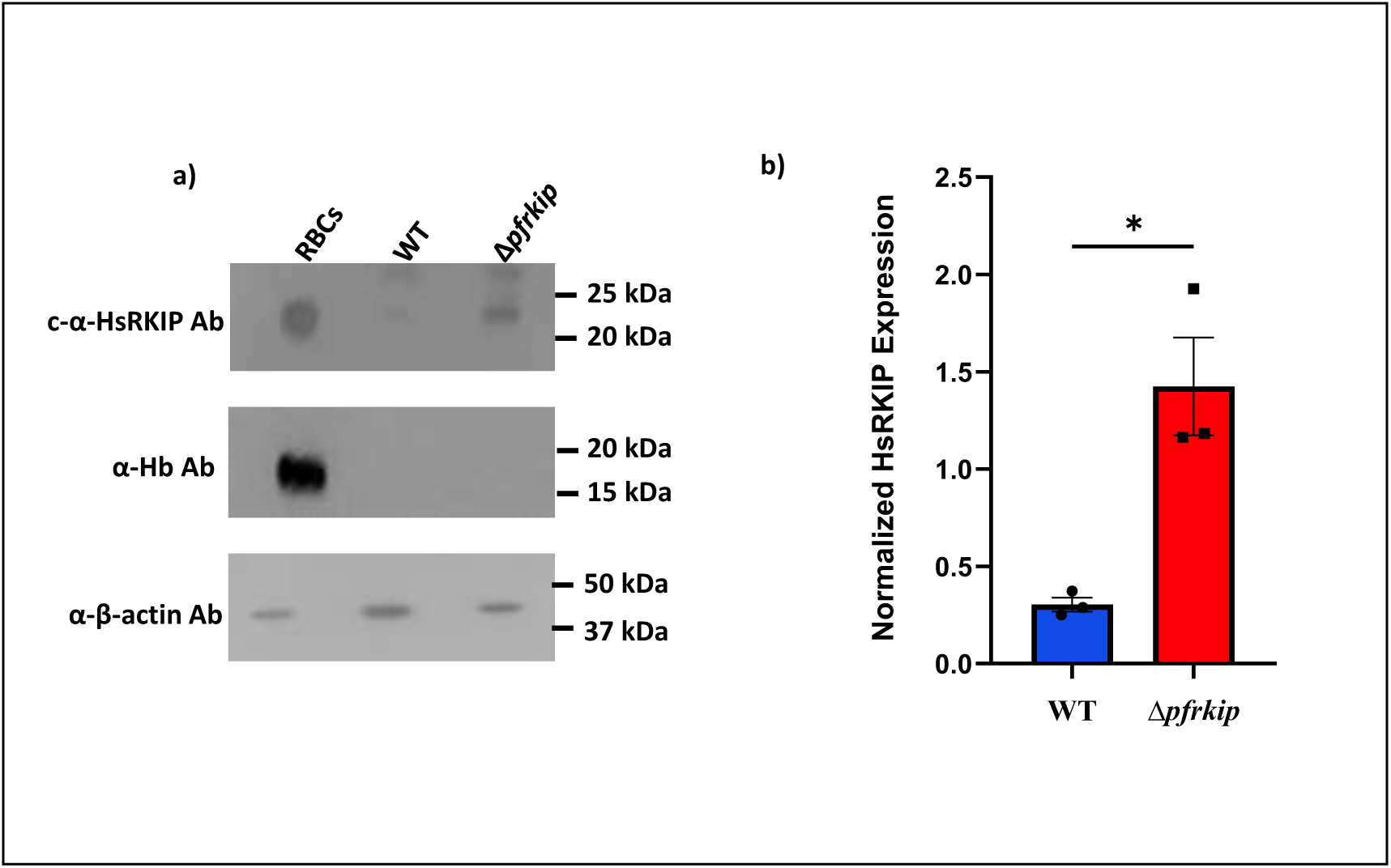
The Δ*pfrkip* parasites show increased import of HsRKIP. **a)** The expression of HsRKIP, hemoglobin (Hb, control), and beta-actin (control) was tested by Western blotting using saponin-lysed, late-stage Δ*pfrkip* and WT parasites. A set of representative blots probed with commercial anti-HsRKIP (c-anti-HsRKIP), anti-Hb and anti-Beta Actin antibody from three independent experiments are shown. RBC lysate is used as a control for the presence of HsRKIP and hemoglobin. **b)** The import of HsRKIP in the Δ*pfrkip* parasites is increased ∼ 5-fold. Bands corresponding to HsRKIP were quantified and normalized to beta-actin and hemoglobin (if detected) using the ImageJ software. The normalized HsRKIP expression on the Y-axis is plotted for the WT and the Δ*pfrkip* parasites on the X-axis. Data represents mean± SEM from three independent biological experiments. The increase in the HsRKIP expression in the Δ*pfrkip* parasites compared to the WT is statistically significant (n=3, *p<0.05, unpaired t-test).

Following validation of antibody specificity and optimization of parasite lysis conditions, we compared HsRKIP levels in WT and *Δpfrkip* parasites. Across multiple independent experiments, we consistently observed a higher abundance of HsRKIP in *Δpfrkip* parasites relative to WT parasites (Fig. 5a). Hemoglobin was undetectable in both parasite samples, indicating complete removal of host RBC cytoplasmic contents and excluding host contamination as a confounding factor (Fig. 5a). Detection of β-actin confirmed equivalent loading of parasite proteins, while RBC lysate served as an additional control (Fig. 5a).

Quantification of HsRKIP signal intensity, normalized to β-actin and hemoglobin (where detectable), revealed an approximately fivefold increase in HsRKIP import in *Δpfrkip* parasites (1.424 ± 0.435) compared to WT parasites (0.3035 ± 0.063) (n = 3; p < 0.05, unpaired t-test) (Fig. 5b). Collectively, these results demonstrate a significant increase in HsRKIP import in *Δpfrkip* parasites, suggesting that enhanced uptake of host HsRKIP may represent an adaptive response to compensate for the loss of PfRKIP.

### Host RKIP interacts with PfCDPK1 within the parasite

We previously demonstrated that PfRKIP interacts with PfCDPK1 within *P. falciparum* parasites^10^. Given the increased import of HsRKIP observed in *Δpfrkip* parasites, we hypothesized that host-derived HsRKIP may functionally compensate for the loss of PfRKIP by engaging PfCDPK1. To test this possibility, we examined whether HsRKIP is phosphorylated by PfCDPK1 and the two proteins interact with each other.

To test whether HsRKIP is phosphorylated by PfCDPK1, we set up an *in vitro* kinase assay with recombinant PfCDPK1 in the presence and absence of calcium and used HsRKIP as an exogenous substrate. After the kinase reaction, the samples were separated on an SDS-PAGE followed by detection of the phosphorylated proteins using ProQ Diamond phosphoprotein stain. The phosphorylation of HsRKIP was increased in the presence of calcium compared to without calcium condition (Fig. S5c) suggesting that HsRKIP is indeed phosphorylated by PfCDPK1.

Next, we tested whether HsRKIP and PfCDPK1 interact with each other in the cellular context. To this end, we generated a transgenic parasite line in the *Δpfrkip* background that overexpresses HsRKIP as a chimeric protein fused at the C-terminus to the promiscuous biotin ligase BioID2^33,34^ and an additional V5 epitope tag (*Δpfrkip^HsRKIPBioID2-V5^*). Firstly, the presence of the HsRKIPBioID2-V5 construct in transfected parasites was confirmed by a diagnostic PCR. Amplification using plasmid-specific primers spanning the cloned region yielded an amplicon of the expected size (1573 bp), whereas control parasites transfected with HsRKIP-V5 construct lacking the BioID2 tag produced an 853 bp amplicon (Fig. S6a).

Expression of HsRKIPBioID2–V5 and HsRKIP-V5 in the respective transgenic parasite lines was verified by Western blot analysis using c-anti-HsRKIP and anti-V5 antibodies. Bands corresponding to the expected molecular weights of HsRKIPBioID2–V5 (∼50 kDa) and HsRKIP-V5 (∼23 kDa) were detected in *Δpfrkip^HsRKIPBioID2–V5^*and *Δpfrkip^HsRKIP-V5^* parasites, respectively, with both antibodies (Fig. S6b and Fig. S6c).

We next assessed global protein biotinylation in the *Δpfrkip^HsRKIPBioID2–V5^*and control, *Δpfrkip^HsRKIP-V5^*, parasites by probing parasite lysates with streptavidin–HRP. As expected, the extent and pattern of biotinylated proteins were markedly higher in *Δpfrkip^HsRKIPBioID2–V5^*parasites compared to the control parasites lacking the BioID2 tag (Fig. 6a). Biotinylated proteins were subsequently affinity-purified from both parasite lines using streptavidin-conjugated magnetic beads. Input samples were probed with anti-V5 antibodies to confirm comparable expression and loading (Fig. 6b, Input).

**Figure 6.**
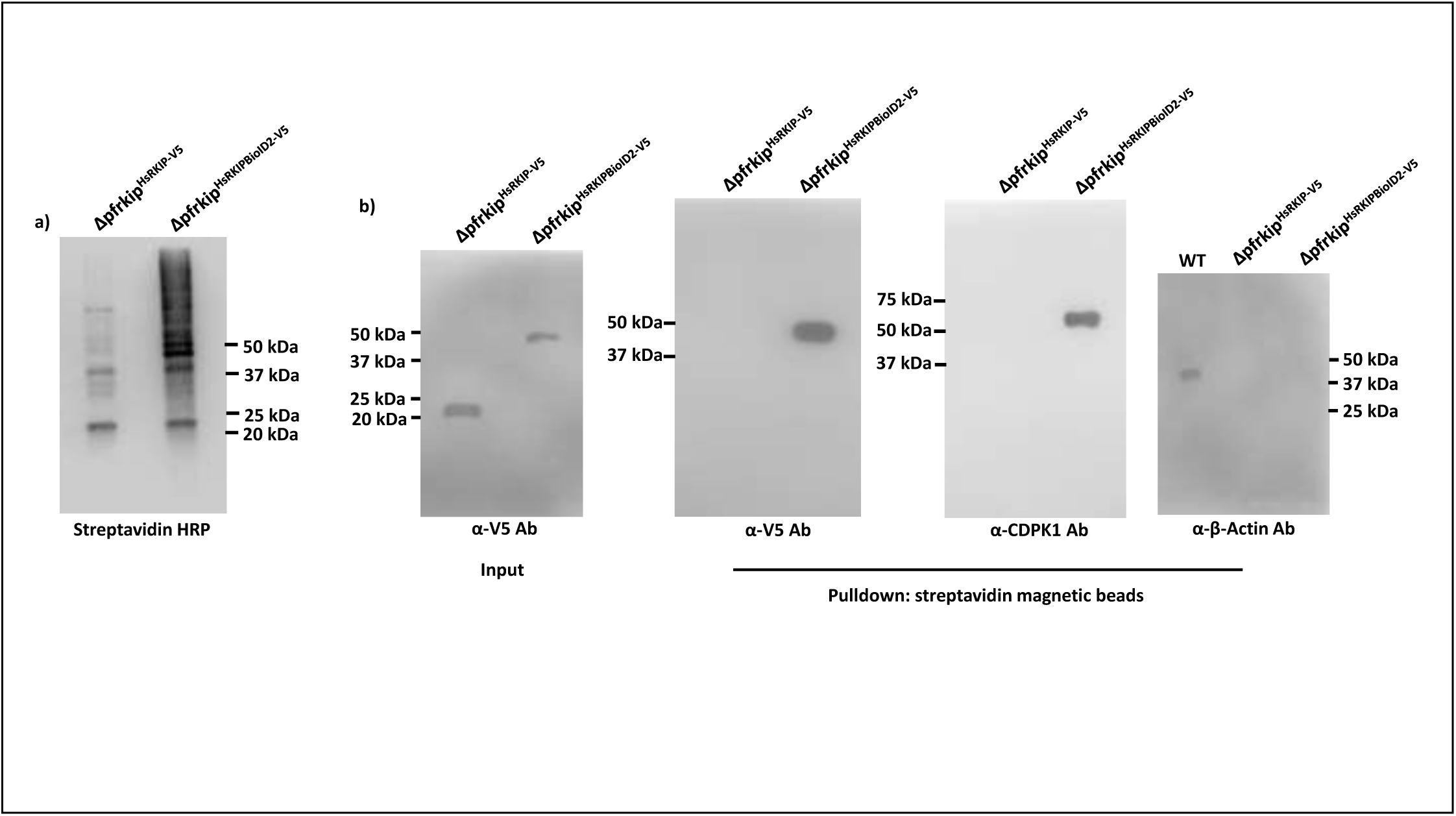
PfCDPK1 and HsRKIP interact in the Δ*pfrkip* parasites. **a)** The Δ*pfrkip* parasites expressing HsRKIPBioID2-V5 (Δ*pfrkip*^HsRKIPBioID2-V5^) show overall increase in the biotinylated proteins compared to the Δ*pfrkip*^HsRKIP-V5^ parasites. The Δ*pfrkip*^HsRKIPBioID2-V5^ and the Δ*pfrkip*^HsRKIP-V5^ parasites were incubated with biotin, harvested, and lysed in RIPA (Radioimmunoprecipitation assay) buffer. The soluble lysates were resolved on the SDS-PAGE, transferred to a PVDF membrane, and probed with streptavidin-HRP antibody to assess the overall biotinylation. Both transgenic parasites show biotinylated proteins. The overall abundance of biotinylated proteins was substantially higher in the HsRKIPBioID2V5-expressing parasites compared to the control. A representative blot from 2 independent biological experiments is shown. **b)** HsRKIP and PfCDPK1 interact within the parasites. The RIPA-lysed soluble fractions from both the transgenic parasites were incubated with streptavidin magnetic beads to pulldown all the biotinylated proteins. The input, lysate before incubation with streptavidin magnetic beads, was probed with anti-V5 antibody to confirm the presence of V5 tagged HsRKIP in the Δ*pfrkip*^HsRKIPBioID2-V5^ and the Δ*pfrkip*^HsRKIP-V5^ parasites. The pull-down fractions were tested for the interaction between PfCDPK1 and HsRKIP. Western blot with anti-V5 antibody confirmed the pulldown of HsRKIPBioID2-V5 (∼ 50 kDa) in the Δ*pfrkip*^HsRKIPBioID2-V5^ parasites, while no HsRKIP-V5 was detected in the control. Western with anti-PfCDPK1 antibody detected a band corresponding to PfCDPK1 (∼ 60 kDa) in the elution fraction of Δ*pfrkip*^HsRKIPBioID2-V5^ parasites, but not in the control parasites. The pull-down samples probed with anti-beta actin antibody show no detection of beta-actin in the Δ*pfrkip*^HsRKIPBioID2-V5^ parasites showing specificity of HsRKIP and PfCDPK1 interaction in the Δ*pfrkip*^HsRKIPBioID2-V5^ parasites. Representative set of Western blots are shown from 2 independent biological experiments. WT-wild type parasite lysate.

The streptavidin pull-down fractions were analyzed by Western blot using anti-PfCDPK1 and anti-V5 antibodies. A band corresponding to HsRKIPBioID2–V5 was detected specifically in pull-downs from *Δpfrkip^HsRKIPBioID2–V5^* parasites (Fig. 6b). Importantly, PfCDPK1 was detected exclusively in the pull-down fraction from *Δpfrkip^HsRKIPBioID2–V5^* parasites and not in the control *Δpfrkip^HsRKIP-V5^* parasites (Fig. 6b), indicating a specific association between HsRKIP and PfCDPK1 inside the parasite. To rule out non-specific protein association, the pull-down samples were probed with anti-β-actin antibody. β-actin was not detected in the pull-down from *Δpfrkip^HsRKIPBioID2–V5^* parasites, whereas it was readily detected in whole-cell lysate used as a control (Fig. 6b).

Collectively, these data demonstrate that host-derived HsRKIP interacts with PfCDPK1 within the parasite, supporting the notion that imported HsRKIP may functionally compensate for the loss of PfRKIP in *Δpfrkip* parasites.

### Host RKIP co-exist with PfCDPK1 in high molecular weight complexes within the parasite

Our results show that PfRKIP and PfCDPK1 form a high molecular weight complex, ranging between 100 kDa and 150 kDa, within the parasite (Fig. 2c). Given that HsRKIP import is increased in *Δpfrkip* parasites and that HsRKIP interacts with PfCDPK1, we hypothesized that, in the absence of PfRKIP, HsRKIP may assemble into high molecular weight complexes analogous to those observed in RKIP-cMyc parasites (Fig. 2c).

To test this, *Δpfrkip^HsRKIP-V5^* schizonts were treated with or without locostatin, followed by covalent cross-linking using DSS. Western blot analysis with the c-anti-HsRKIP antibody revealed a prominent high molecular weight complex between 100 kDa and 150 kDa in the locostatin-treated parasites (Fig. 7a, black arrow). The corresponding complex was also detected with anti-PfCDPK1 antibody (Fig. 7a, black dotted arrow), indicating co-association of HsRKIP and PfCDPK1. Additionally, a higher molecular weight complex exceeding 150 kDa was observed with both antibodies (white solid/dotted arrows), suggesting the coexistence of HsRKIP and PfCDPK1 in larger assemblies. Notably, this >150 kDa complex was not detected with anti-PfRKIP antibody in PfRKIP-cMyc parasites (Fig. 2c) but was observed with anti-HsRKIP antibody (Fig. 7a), indicating that, unlike PfRKIP, HsRKIP can participate in higher molecular weight complexes with PfCDPK1. Anti-β-actin antibodies confirmed equal loading across all samples (Fig. 7a).

**Figure 7.**
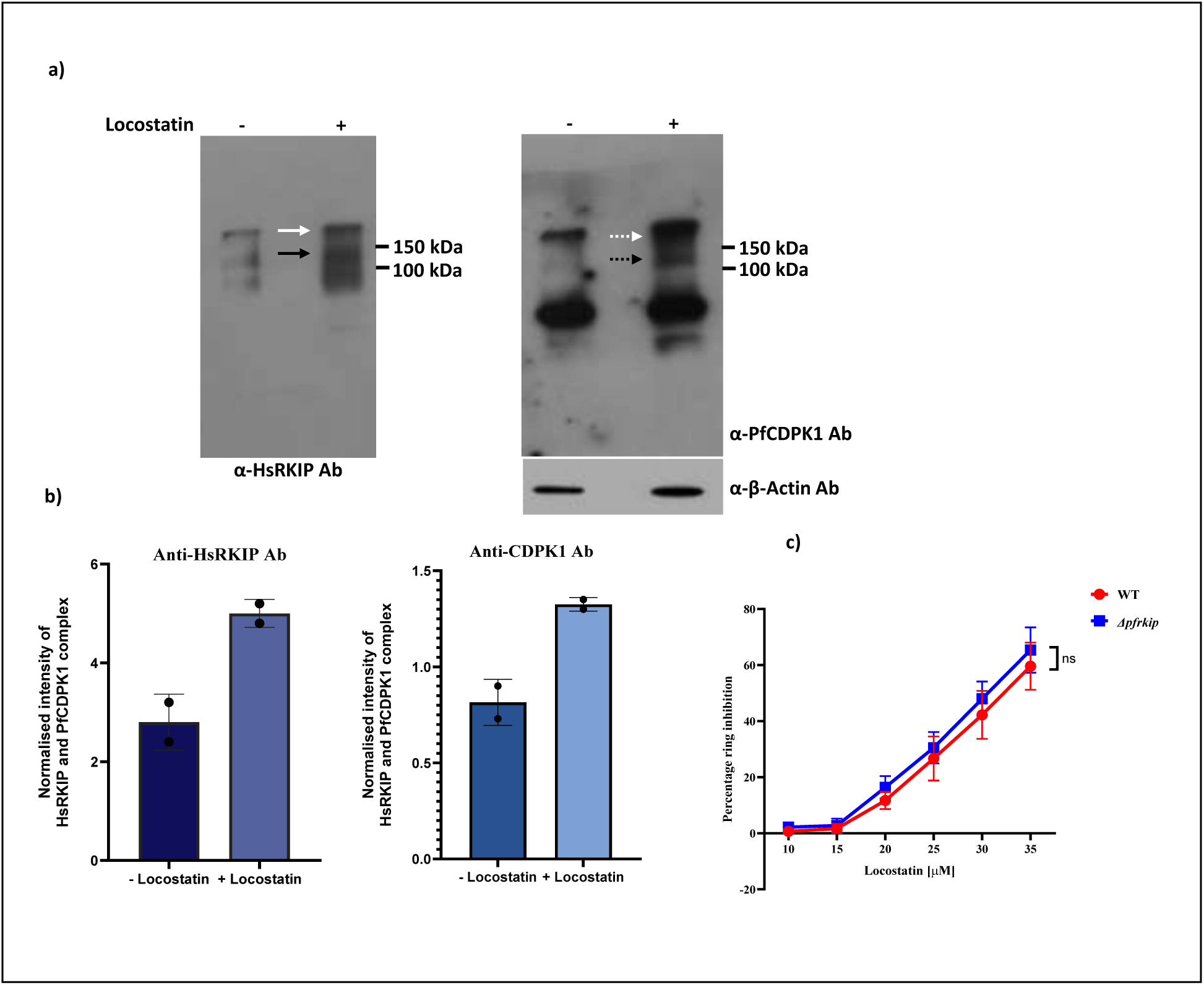
Locostatin enhances the interaction between PfCDPK1 and HsRKIP and show similar sensitivity in the Δ*pfrkip* parasites as the WT. a) Late-stage parasites treated with locostatin were crosslinked with disuccinimidyl suberate (DSS), resolved on the SDS-PAGE, and transferred onto a PVDF membrane. The membrane was probed with anti-HsRKIP and anti-PfCDPK1 antibodies to test the presence of protein complex between PfCDPK1 and HsRKIP. In the presence of locostatin, a high molecular weight complex (between 100 kDa and 150 kDa) is detected with anti-HsRKIP (black arrow) and anti-PfCDPK1 antibodies (dotted black arrow), suggesting enhanced interaction between PfCDPK1 and HsRKIP with locostatin treatment. A higher molecular weight complex, > 150 kDa is also observed with the anti-HsRKIP (white arrow) and anti-PfCDPK1 antibodies (dotted white arrow). A parallel blot probed with anti-β-actin antibody shows presence of parasite material in the two samples. Represented blots from two independent experiments are shown. **b)** The intensity of the high molecular weight complex comprising of HsRKIP and PfCDPK1 increases in the presence of locostatin. The intensity of the high molecular weight complex (between 100 kDa and 150 kDa) in the presence and absence of locostatin detected with the anti-HsRKIP and anti-PfCDPK1 antibodies was normalized to β-actin. The histograms show an approximately 1.8-fold and 1.6-fold increase in the intensity of the complex in the presence of locostatin with the anti-HsRKIP and anti-PfCDPK1 antibodies, respectively. These results confirm enhancement in the interaction between PfCDPK1 and HsRKIP in the presence of locostatin. The error bars represent standard error of the mean from 2 independent biological experiments. **c)** Highly synchronized schizonts of Δ*pfrkip* and WT parasites were treated with varying concentrations of locostatin. The percent ring parasitemia was estimated after 12-14 h through FACS. Locostatin treatment leads to a dose-dependent decrease in the proportion of infected RBCs. The graph shows the percent decrease in ring parasitemia in the WT and the Δ*pfrkip* parasites on the Y-axis at increasing locostatin concentrations [in μM] on the X-axis. Locostatin inhibits both WT and Δ*pfrkip* parasites; however, the Δ*pfrkip* parasites exhibit modestly higher sensitivity to locostatin, as evidenced by a slightly higher inhibition than WT parasites across all tested concentrations. The difference is not statistically significant (n=3, p>0.05, 2way ANOVA, Sidak’s multiple comparison test). Data are expressed as mean ± SEM (n= 3 biological experiments).

Densitometric analysis of the complexes between 100 kDa and 150 kDa, normalized to β-actin, revealed that the mean intensity in the presence versus absence of locostatin was 5.0 ± 0.2 and 2.8 ± 0.4, respectively, with the c-anti-HsRKIP antibody, and 1.325 ± 0.025 and 0.815 ± 0.085, respectively, with the anti-PfCDPK1 antibody (Fig. 7b).

These findings suggest that, in the absence of PfRKIP, imported HsRKIP assembles into high molecular weight complexes with PfCDPK1, potentially compensating for the loss of PfRKIP function in *Δpfrkip* parasites (Fig. 7).

### Δ*pfrkip* parasites show similar sensitivity to locostatin as the WT

Our results suggest that locostatin inhibits the growth of WT parasites through sequestration of PfCDPK1 in complex with PfRKIP. In the Δ*pfrkip* parasites, the import of the host RKIP is 5-fold increased and it interacts with PfCDPK1 just like the PfRKIP. Moreover, in the presence of locostatin host RKIP forms similar size complex with PfCDPK1 as in the WT parasites. Based on these findings, we hypothesized that locostatin may have a similar inhibitory effect on the Δ*pfrkip* parasites as in the WT. To this end, synchronized mature schizonts (38–42 HPI) of WT and *Δpfrkip* parasites were exposed to increasing concentrations of locostatin, and the formation of ring-stage parasites was assessed after 12-14 h, in the subsequent cycle, through fluorescence activated cell sorter (FACS). Treatment with locostatin resulted in a dose-dependent reduction in the growth of both parasites (Fig. 7c). Notably, the percent inhibition in parasite growth was moderately higher in the *Δpfrkip* parasites than the WT at all the concentrations of locostatin tested. However, the difference was not statistically significant (Fig. 7c; n=3; p>0.05, 2way ANOVA, Sidak’s multiple comparison test). The samples used for the FACS were also read through SYBR Green I-assay. SYBR Green I-assay result showed a similar concentration-dependent decrease in parasite-derived fluorescence, consistent with reduction in the parasite proliferation (Fig. S7). Furthermore, a modest, non-significant increase (n=3, p>0.05, 2way ANOVA, Sidak’s multiple comparison test) in sensitivity to locostatin was observed in the *Δpfrkip* parasites compared to the WT in the SYBR Green I assay similar to the result with the FACS (Fig. 7c and Fig S7). Comparable sensitivity of *Δpfrkip* parasites to locostatin treatment suggests that PfCDPK1 is sequestered by the locostatin-modified host RKIP. This is analogous to the sequestration of PfCDPK1 by PfRKIP upon locostatin treatment of the WT parasites.

## METHODS

### *In vitro* culture of malaria parasite

NF54 strain of *Plasmodium falciparum* was grown under *in vitro* conditions in O+ human red blood cells (Rotary Blood Bank, New Delhi) at 2 - 4 % hematocrit. RPMI1640 medium containing L-glutamine and supplemented with 25 mM HEPES (Sigma-Aldrich), 50 µg/mL Hypoxanthine (Sigma-Aldrich), 0.5 % Albumax I (ThermoFisher Scientific), 25 mM sodium bicarbonate (Sigma-Aldrich) and 10 μg/mL gentamycin (ThermoFisher Scientific) was used as described earlier^35^. The parasites were propagated at 37 °C under a gaseous environment of 5 % O_2_, 5 % CO_2_, and 90 % N_2_.

### Generation of *pfrkip* knock-out parasites

Transgenic parasites with complete knock-out of endogenous *pfrkip* gene were generated through CRISPR/Cas9. A 20-nucleotide guide sequence (5’ TTTTAGATATTGCTGGAACG 3’), corresponding to 134 to 153 bp within the open reading frame of *pfrkip* adjacent to the protospacer adjacent motif (PAM; 5’ GGG 3’), was inserted into the pL6Cas9 plasmid with In-Fusion (Takara Bio), using the primer pair GRKIPKOF/GRKIPKOR, generating pL6Cas9-gRKIP plasmid. The 5’ and 3’ homology arms of 380 and 267 base pairs were amplified using primer pairs, 5’HRRKIPKOF/5’HRRKIPKOR and 3’HRRKIPKOF/3’HRRKIPKOR, respectively, from the genomic DNA and cloned into SpeI/AflII and EcoRI/NcoI restriction sites, respectively in the pL6Cas9-gRKIP plasmid. The final plasmid, *pL6-Cas9-pfrkipko* was transfected into the ring-stage parasites as previously described^36^. The transfected parasites were selected with 2.0 nM WR99210. Drug-resistant parasites appear after 3 weeks of transfection. Clonal parasites were obtained through limiting dilution.

### RNA isolation and validation of *rkip* knockout

Schizont-stage parasites (42–48 hours post-invasion, HPI) from wild-type (WT) and RKIP knockout (*Δpfrkip*) strains were harvested and processed for total RNA extraction as described elsewhere^37^ using the RNeasy Mini Kit (Qiagen), following the manufacturer’s instructions. To ensure complete removal of genomic DNA contamination, RNA samples were subjected to two DNase treatments: an on-column DNase I digestion during RNA purification and a second post-extraction treatment using the TURBO DNA-free™ Kit (ThermoFisher Scientific). The quality and concentration of purified RNA were assessed using a NanoDrop spectrophotometer (ThermoFisher Scientific). First-strand cDNA was synthesized from purified RNA using the PrimeScript™ 1st Strand cDNA Synthesis Kit (Takara Bio), according to the manufacturer’s protocol. The synthesized cDNA was used to assess the presence or absence of *rkip* transcripts in WT and *Δpfrkip* parasites by PCR amplification using the primer sets RKIPFRT/RKIPRRT, RKIPS96A/PfRKIPG4T1_R, and RKIPV5F1/PfRKIPG4T1_R. Genomic DNA contamination was further evaluated by PCR amplification of *pfcdpk1* using cDNA as a template. The amplification yielded a smaller product compared to genomic DNA due to intron exclusion, confirming the absence of contaminating genomic DNA in the cDNA samples.

### Gene expression analysis in WT and *Δpfrkip* parasites

cDNA synthesized from WT and *Δpfrkip* parasites was used to assess the expression of *pfcdpk1* and *PF3D7_0303900* by PCR amplification using gene-specific primer sets, CDPK1RTF/CDPK1RTR and PEBP3900FP/PEBP3900RP. A house-keeping gene, glyceraldehyde phosphate dehydrogenase (*gapdh*) was amplified using the primer pair, GAPDHRTF/GAPDHRTR as an internal control. Gene expression levels were quantified following normalization with the housekeeping gene *pfgapdh*, which served as an internal control.

### Generation of HsRKIP overexpressing parasite strain

To study the interaction of HsRKIP with PfCDPK1, we used the proximity biotinylation strategy^38–40^. The full-length HsRKIP was fused at the C-terminus with a modified biotin ligase, BioID2^33,34^ and expressed episomally in pDC2 plasmid. For generating the final construct in pDC2 plasmid, BioID2 was amplified from pm2gt-hsp101-bioid2^34^ plasmid using the primer pair, pET42BIOID2F/pET42BIOID2R and cloned between XhoI and AvrII sites in pET42a(+) plasmid containing HsRKIP^10^. The double digestion positive clones were sequence verified and used to amplify the complete HsRKIP-BioID2 sequence using the primer pair, pDC2AHsRKIPFWD/pDC2RKIPBIOID2_R. Simultaneously, the HsRKIP gene alone was amplified using another primer set, pDC2AHsRKIPFWD/pDC2AHsRKIPREV. The amplicons were inserted into pDC2 expression plasmid at the AvrII and MluI restriction sites followed by the transformation of the XL 10-Gold competent cells. Successful integration of the gene segments was verified by double digestion followed by DNA sequencing. The sequence-confirmed plasmids were purified using NucleoBond Xtra Maxi EF (Macherey-Nagel). The Δ*pfrkip* parasites were transfected with the purified plasmids to obtain the transgenic *Δpfrkip^HsRKIPBioID2-V5^* and *Δpfrkip^HsRKIP-V5^* parasites.

### Locostatin mediated inhibition of parasite growth

Highly synchronized *P*. *falciparum* schizonts (38-42 HPI) were adjusted to a final parasitemia of 0.5 % at 2 % haematocrit and were incubated with different concentrations of locostatin (10 μM, 15 μM, 20 μM, 25 μM, 30 μM, and 35 μM) in a 24-well plate. The newly formed rings and the unruptured schizonts after 10 - 12 h of treatment were then scored by Giemsa staining under a light microscope. The percentage ring or schizont parasitemia on the Y-axis was plotted against different concentrations of locostatin (μM) on the X-axis from 3 independent biological experiments in duplicate. Statistical significance was calculated using one-way ANOVA in GraphPad Prism 10.6.1.

We also employed an automated SYBR Green I-based fluorescence assay to test the effect of locostatin on the growth of the WT parasites. The experiment was set-up as described above. Aliquots (100 μl) of the WT and uninfected RBC controls were collected after 10 - 12 h of locostatin treatment and stored at -80 °C for subsequent analysis. The samples were thawed and lysed in equal volume of 2x lysis buffer [20 mM Tris (pH 7.5), 5 mM EDTA, and 0.008 % (wt/vol) saponin], followed by incubation with 5X SYBR Green I for 30 min at 37 °C. Fluorescence was measured using a Fluostar Optima plate reader (BMG Labtech) with excitation and emission wavelengths set at 485 nm and 520 nm, respectively, with a gain of 1000^41,42^. Three independent experiments were performed in duplicate. The data was plotted with fluorescence intensity on the Y-axis, indicating parasite growth, against the concentration of locostatin on the X-axis. The graph was prepared using the GraphPad Prism version 10.6.1. Statistical test (RM One-Way ANOVA) was performed on the data obtained from the three independent biological experiments.

### Effect of locostatin on the interaction of recombinant PfRKIP and PfCDPK1

ELISA was also used to test the effect of locostatin on the interaction of PfCDPK1 and PfRKIP. Locostatin is known to modify mammalian RKIP through alkylation of a histidine residue^29^. For conditions requiring testing the effect of modified PfRKIP on interaction with PfCDPK1, the recombinant PfRKIP was pre-incubated with or without different concentrations of locostatin: 50 μM, 100 μM, 200 μM, 400 μM, and 800 μM at 16 °C for 2 h before adding to PfCDPK1 coated wells. The composition of the interaction buffer is 50 mM HEPES (pH 8.0), 250 mM potassium acetate, 5 mM magnesium acetate, 150 mM NaCl. To test the effect of locostatin on the interaction of PfCDPK1 and unmodified PfRKIP, the recombinant PfRKIP was not pre-incubated with locostatin rather locostatin was added concomitantly along with other reagents in the interaction buffer. All the experiments were performed in triplicate and mean ± SEM was calculated. The OD_492_ on the Y-axis representing binding of either modified or unmodified PfRKIP with PfCDPK1 in presence and absence of locostatin was plotted with locostatin concentration on the X-axis. Graphs were plotted using GraphPad Prism version 10.6.1.

### Effect of phosphorylation on the interaction of PfCDPK1 with locostatin-modified PfRKIP

To test the effect of phosphorylation on the interaction between PfCDPK1 and chemically modified PfRKIP, we used an ELISA-based assay. The recombinant PfCDPK1 was immobilized in a 96-well plate as previously described^10^. PfRKIP taken at different concentrations (21.87, 43.75, 87.5, 175, 350, and 700 nM) was alkylated with 800 µM locostatin at 16 °C for 2 h in the interaction buffer mentioned above. DMSO was employed as a vehicle control in cases where no modification of PfRKIP was required.

Alkylated PfRKIP was incubated with PfCDPK1 in a 96-well plate, either in the presence or absence of 100 µM ATP to assess the effect of phosphorylation on the interaction between the two proteins. Additionally, 10 mM CaCl₂ was added to the reaction buffer to activate PfCDPK1. Post-incubation, the binding of modified PfRKIP was detected using anti-PfRKIP antibodies, followed by an anti-rat HRP-conjugated secondary antibody (1:35,000). The enzymatic reaction was initiated by addition of o-phenylenediamine dihydrochloride (OPD) substrate and hydrogen peroxide. The reaction was allowed to proceed for 30 min at RT followed by the addition of sulphuric acid to stop the reaction. The plate was read at 492 nm using a Multiskan SkyHigh spectrophotometer (ThermoFisher Scientific). The absorbance values at 492 nm (Y-axis) were plotted against the concentrations of recombinant PfRKIP modified by pre-treatment with locostatin (X-axis). Statistical analysis was performed using a multiple paired t test in GraphPad Prism 10.6.1, based on three independent biological replicates, each performed in duplicate.

### Effect of locostatin on the interaction between PfRKIP and PfCDPK1 within parasites

To test the effect of locostatin on the interaction between PfCDPK1 and PfRKIP in *P. falciparum*, a parasite line overexpressing PfRKIP, RKIP-c-Myc^10^ was used. The parasites were synchronized by sorbitol treatment, followed by isolation of late-stage trophozoites and schizonts using Percoll/sorbitol gradient. Schizont-stage parasites were incubated with fresh RBCs for 4 h, then re-synchronized using 5 % sorbitol to obtain 0-4 h ring-stage parasites. The parasites were cultured until 32-36 h trophozoite stage followed by purification using Percoll/sorbitol gradient. The 32-36 h trophozoites were divided into two groups: one without locostatin treatment (control) and another treated with 30 µM locostatin in 1 mL cRPMI in a 24-well plate. The 30 µM locostatin concentration was chosen as it shows non-significant and negligible effect on the egress of the parasites from the infected RBCs. Both the treatment groups were incubated for an additional 12 h at the optimal parasite growth conditions as described in the parasite maintenance section. After reaching the 44-48 h schizont stage, the parasites were pelleted by centrifugation at 6000 rpm for 5 min, washed twice with 1X PBS, and incubated for 30 min at RT with 1.5 mM disuccinimidyl suberate (DSS), a cell permeable crosslinking agent. The crosslinking reaction was terminated by the addition of 20 mM Tris-HCl (pH 7.6) for 15 min at RT. The cross-linked parasite pellet was washed twice with 1× PBS containing 0.01 % saponin. The parasite pellets were then resuspended in 1X SDS sample buffer and separated by SDS-PAGE. The proteins were transferred onto a PVDF membrane followed by blocking with 5 % skim milk prepared in 1XPBS. The membrane was probed with anti-CDPK1 (1:2000) or anti-PfRKIP (1:2000) antibody. Following incubation with the primary antibodies, the membrane was incubated with the compatible secondary antibodies at 1:5000 dilution, anti-rabbit HRP-conjugated for the PfCDPK1 and anti-rat HRP-conjugated for PfRKIP. Signals were detected using enhanced chemiluminescence (ECL) followed by exposure on to the X-ray film.

### Effect of locostatin on the interaction between HsRKIP and PfCDPK1 within *Δpfrkip* parasites

To test the effect of locostatin on the interaction between PfCDPK1 and HsRKIP in the parasite a genetically modified parasite line overexpressing HsRKIP in the *Δpfrkip* background was used. The experiment was set-up in the same way as described above for the ‘effect of locostatin on the interaction between PfRKIP and PfCDPK1 within parasites’. The c-anti-HsRKIP antibody was used to detect HsRKIP instead of anti-PfRKIP antibody. Anti-rabbit HRP-conjugated secondary antibody was used at 1:5000 dilution.

### Proximity biotinylation to study the interaction of HsRKIP with PfCDPK1 within the parasites

Late-stage trophozoite/schizont-stage parasites expressing HsRKIPBioID2-V5 or HsRKIP-V5 alone (control) were incubated with 200 µM biotin for 48 h to allow biotinylation of proteins. The culture media containing biotin was replenished every 24 h. After the incubation period, the parasites were harvested by centrifugation at 900 g for 3 min, and the supernatant was carefully removed. The RBC pellet containing parasites was treated with 0.01 % saponin in 1X PBS to selectively lyse RBCs, indicated by a dark brown parasite pellet. The saponin-treated pellet was then washed twice with 1X PBS, and the parasite pellets were stored at -80 °C until further use.

To pull-down all the biotinylated proteins, the *Δpfrkip^HsRKIPBioID2-V5^*and the *Δpfrkip^HsRKIP-V5^* control parasites were lysed in RIPA buffer and incubated on ice for 1 h. The lysates were cleared by centrifugation at 12,000 rpm for 30 min at 4 °C, and the soluble fraction was collected in a separate tube. The lysates were diluted 2.5-fold with the resuspension solution: 50 mM Tris-Cl (pH 7.5) and 100 mM NaCl. Streptavidin magnetic beads were used to capture the biotinylated proteins following the manufacturer’s protocol. The diluted lysates were incubated with the streptavidin beads overnight at 4 °C with constant agitation. Following incubation, the beads were pelleted, and the flow-through was discarded. The beads were washed twice with TBST (0.01 % Tween-20) to remove non-specifically bound proteins. The biotinylated proteins were released from the beads by resuspending them in 100 µl of 1X SDS sample buffer.

The samples were separated on SDS-PAGE and transferred to a PVDF membrane. The membrane was probed with specific primary antibodies against V5, PfCDPK1, and β-actin, followed by appropriate secondary antibodies, as per the previously described Western blot protocol. Protein detection was carried out using enhanced chemiluminescence (ECL), and the signal was detected by exposure on to the X-ray film.

### Effect of Locostatin-mediated, PfRKIP-PfCDPK1 complex formation on the transphosphorylation activity of PfCDPK1

To evaluate the impact of locostatin-mediated, PfRKIP-PfCDPK1 complex formation on the trans-phosphorylation activity of PfCDPK1, an *in vitro* kinase was performed as described previously^10^. Briefly, 5 µg of PfRKIP was incubated with or without 800 µM locostatin in an interaction buffer of the composition: 50 mM HEPES (pH 8.0), 250 mM potassium acetate, 5 mM magnesium acetate, 150 mM NaCl at 16 °C for 2 h. Following the pre-incubation, 10 mM ATP, 5 µg myelin basic protein (MBP), 250 ng PfCDPK1, and 5 mM CaCl₂ were added to initiate the kinase reaction. The kinase assay was allowed to proceed at 30 °C for 1 h. As a control, an additional reaction was set up where PfCDPK1 was incubated with 10 mM ATP, 5 µg MBP, with and without 800 µM locostatin in the absence of PfRKIP to assess the direct effect of locostatin on PfCDPK1’s kinase activity under identical conditions.

The reactions were stopped by adding SDS sample buffer, and 25 µL of each sample was loaded onto an SDS-PAGE gel. The gel was subsequently stained using the Pro-Q Diamond phosphoprotein staining solution as per the manufacturer’s instructions. Phosphorylated MBP was detected using a GelDoc imaging system under UV light, and the intensities of the phosphorylated MBP bands were quantified using ImageJ software version 1.53m. Normalized phosphorylation of MBP was plotted for the with and without locostatin conditions using GraphPad Prism version 10.6.1.

### Comparison of the asexual growth of Δ*pfrkip* with the WT parasite using bright-field microscopy and SYBR Green I

The asexual growth of *Δpfrkip* was compared with the WT parasites using Giemsa staining and SYBR Green I staining. The *Δpfrkip* and the WT parasites were synchronized with two consecutive cycles of sorbitol treatment. The trophozoite stage parasites were seeded at 0.2 % parasitemia at 2 % hematocrit in a 24-well plate. Aliquots of both the parasites were drawn after every 24 h for 4 days and smeared on a glass slide for Giemsa counting. Around 2500 total RBCs were counted in duplicate for each slide for each parasite strain. Parasitemia for each day was plotted on the Y-axis against the days on the X-axis.

We also employed an automated SYBR Green I-based fluorescence assay to compare the asexual growth of the *Δpfrkip* and the WT parasites. The experiment was set-up as described above. Aliquots of the *Δpfrkip*, WT, and uninfected RBC controls were collected for 4 consecutive days and stored at -80 °C for subsequent analysis. The samples were thawed and lysed in equal volume of 2x lysis buffer [20 mM Tris (pH 7.5), 5 mM EDTA, and 0.008 % (wt/vol) saponin], followed by incubation with 5X SYBR Green I for 30 min at 37 °C. Fluorescence was measured using a Fluostar Optima plate reader (BMG Labtech) with excitation and emission wavelengths set at 485 nm and 520 nm, respectively, with a gain of 1000^41,42^. Two independent experiments were performed in duplicate. The data was plotted with fluorescence intensity on the Y-axis, indicating parasite growth, against the days on the X-axis. The graph was prepared using the GraphPad Prism version 10.6.1.

### Bright-field microscopy and SYBR Green I assay to test the effect of locostatin on the WT parasites

Highly synchronized 38-42 HPI schizonts of WT parasites were seeded at 0.5 % parasitemia in a 24-well plate with a final volume of 500 μL. These parasites were then treated with various concentrations of locostatin: 10 μM, 15 μM, 20 μM, 25 μM, 30 μM, and 35 μM. Samples were collected 12-14 h post-treatment, and the parasite growth was determined using bright-field microscopy as described above.

We also employed an automated SYBR Green I-based fluorescence assay to test the effect of locostatin on the growth of the WT parasites. The experiment was set-up as described above. Aliquots (100 μl) of the WT and uninfected RBC controls were collected after 10 - 12 h of locostatin treatment and stored at -80 °C for subsequent analysis. The samples were thawed and lysed in equal volume of 2x lysis buffer [20 mM Tris (pH 7.5), 5 mM EDTA, and 0.008 % (wt/vol) saponin], followed by incubation with 5X SYBR Green I for 30 min at 37 °C. Fluorescence was measured using a Fluostar Optima plate reader (BMG Labtech) with excitation and emission wavelengths set at 485 nm and 520 nm, respectively, with a gain of 1000 ^43,44^. Three independent experiments were performed in duplicate. The data was plotted with fluorescence intensity on the Y-axis, indicating parasite growth, against the concentration of locostatin on the X-axis. The graph was prepared using the GraphPad Prism version 10.6.1. Statistical test (RM One-Way ANOVA) was performed on the data obtained from the three independent biological experiments.

### Detection of HsRKIP in RBC and specificity of commercial HsRKIP antibody

The detection of HsRKIP in RBCs was performed through Western blot analysis. To prepare the RBC lysate, 10 µL of packed RBCs was boiled in 1X reducing SDS sample buffer. Additionally, 50 µg of purified recombinant HsRKIP protein was run as a positive control, while 50 µg of PfRKIP protein was used as a negative control to verify the specificity of the antibody. All samples were resolved on a 12 % SDS-PAGE gel and transferred to a PVDF membrane. The membrane was blocked and incubated with a commercial anti-HsRKIP (c-α-HsRKIP) antibody (G-Biosciences) at a dilution of 1:5000, followed by washing and subsequent incubation with a secondary anti-rabbit HRP-conjugated antibody (1:5000). The blot was developed using an enhanced chemiluminescence (ECL) substrate, and the signal was visualized by exposing the membrane onto a X-ray film.

### Western blot analysis to compare the level of HsRKIP in the WT and the Δ*pfrkip* parasites

For the detection of HsRKIP in the late-stages, highly synchronized *P. falciparum* parasites were isolated using the Percoll-sorbitol gradient method. The synchronized trophozoite and schizont-stage parasites were split into two equal halves. One was treated with 0.01 % saponin in 1X PBS to lyse the RBCs, followed by repeated washing with 1X PBS at 6000 rpm for 5 min until the parasite pellet turned dark brown, indicating successful RBC lysis. The other half was washed three times with 1X PBS without saponin treatment, keeping the RBCs intact. Both the samples were then incubated with 1X SDS reducing buffer on ice for 1 h, followed by heating at 95 °C for 5 min. The boiled samples were centrifuged at 12,000 rpm for 30 min at 4 °C. The supernatants were collected in separate tubes and separated on a SDS-PAGE followed by transfer to a PVDF membrane. The membrane was blocked with 5 % skim milk in TBST (0.05 % Tween-20) for 1 h at RT to prevent nonspecific binding. The blot was then incubated with c-α-HsRKIP antibody at a dilution of 1:1000 for overnight at 4 °C. After washing the membrane with TBST, it was incubated with a secondary HRP-conjugated anti-rabbit antibody (1:5000 dilution) for 1 h at RT. The bound secondary antibody was detected using enhanced chemiluminescence (ECL), and the signal was visualized by exposure on to an X-ray film.

### Flow cytometric and SYBR Green I assay to test the effect of locostatin on the *Δpfrkip* parasites

Highly synchronized 38-42 HPI schizonts of *Δpfrkip* parasites were seeded at 0.5 % parasitemia in a 24-well plate with a final volume of 500 μL. These parasites were then treated with various concentrations of locostatin: 10 μM, 15 μM, 20 μM, 25 μM, 30 μM, and 35 μM. Samples were collected 12-14 h post-treatment, and the percentage parasitemia of the ring-infected RBCs was determined using flow cytometry as previously described ^44,45^. In brief, 20 μL of the parasite culture from both strains was first washed with 1X phosphate-buffered saline (PBS) and then resuspended in a final volume of 380 μL of 1X PBS. The samples were incubated with 5X SYBR Green I solution for 30 min at RT in dark. Following incubation, the samples were centrifuged at 500 × g for 5 min, washed twice with 1X PBS, and resuspended in 500 μL of 1X PBS. Parasitemia was quantified using a BD FACSCalibur flow cytometer. Samples from each day were also analysed through SYBR Green I fluorescence assay as described above.

Data acquisition via flow cytometry involved collecting 100,000 events per sample, with initial gating based on unstained uninfected red blood cells (RBCs) to assess background autofluorescence. The acquired data was analyzed using CellQuest Pro software version.

## DISCUSSION

PfRKIP interacts with PfCDPK1, and the two proteins preferentially co-localize at the apical end of the merozoite^10^. PfCDPK1 possesses specific N-terminal motifs that facilitate its anchoring to the parasite membrane^46^, while PfRKIP binds phosphorylated phosphatidylinositols, which may be critical for its spatial localization at the membrane, thereby enabling its interaction with PfCDPK1. Using locostatin, a pharmacological inhibitor of mammalian RKIP^30^, we found that parasite invasion into RBCs is significantly impaired. Locostatin irreversibly alkylates His86 in mammalian RKIP, reducing its interaction with Raf-1 kinase and GRK2, ultimately modulating MAPK signaling and cell proliferation^29,47^. In some contexts, locostatin-modified RKIP retains interactions with specific partners while losing others, suggesting a differential regulatory effect of alkylation on protein-protein interactions ^29,48^. Remarkably, in contrast to mammalian RKIP, pre-treatment of PfRKIP with locostatin enhanced its interaction with PfCDPK1, whereas simultaneous addition of locostatin did not affect the interaction. His86 in HsRKIP is conserved as His92 in PfRKIP, and this residue is critical for PfRKIP-PfCDPK1 interaction, as its substitution with alanine diminishes binding^10^. We propose that locostatin-mediated alkylation of His92 may stabilize additional contacts with PfCDPK1, increasing binding affinity.

Pre-treatment of PfRKIP with locostatin enhances its sequestration of PfCDPK1, reducing the kinase’s availability to phosphorylate exogenous substrates. DSS-mediated cross-linking revealed the formation of a high molecular weight complex (100–150 kDa) containing PfRKIP and PfCDPK1. The molecular weight of the complex exceeds the sum of the individual proteins, suggesting incorporation of additional parasite factors^49^. The presence of such a complex in mature schizonts, and potentially in free merozoites, implies a regulatory role in sequential phosphorylation events necessary for RBC invasion. Notably, we observed a distinct PfCDPK1-containing complex of >150 kDa, devoid of PfRKIP, whose intensity decreased upon locostatin treatment. This suggests that locostatin “locks” PfCDPK1 into a PfRKIP-bound complex, precluding its assembly into higher-order complexes with other invasion-related proteins such as 14.3.3l and PKAr^49,50^. Collectively, these observations indicate that PfRKIP may act as a scaffold, orchestrating the formation of intermediate complexes critical for the assembly of invasion-competent molecular machinery.

Whole-genome mutagenesis has suggested an essential role for PfRKIP in the asexual cycle^31^. Contrary to this prediction, CRISPR/Cas9-mediated disruption of the PfRKIP revealed no significant defect in parasite growth under *in vitro* culture conditions. These findings indicate that PfRKIP is not strictly essential under culture conditions, or that compensatory mechanisms may exist. Transcript analysis of another PEBP domain-containing protein, Pf3D7_0303900, and *pfcdpk1* revealed comparable expression in *Δpfrkip* and WT parasites, suggesting no transcriptional compensation by these genes.

Human RBCs contain abundant HsRKIP, which is imported by *P. falciparum* during its development^26–28^. Subcellular fractionation, mass spectrometry, and immunofluorescence studies confirm that HsRKIP localizes within the parasite cytoplasm, independent of host cell contamination^26–28^. Notably, *Δpfrkip* parasites exhibited a ∼5-fold increase in HsRKIP import relative to WT parasites, suggesting a potential compensatory mechanism to restore PfRKIP function. RKIP has been suggested to act as a “sponge” to sequester locostatin in mammalian systems^51^. It means that the effective intracellular concentration of locostatin available for PfRKIP modification in the WT parasites is likely lower than the estimated values. Therefore, it would require higher concentrations of locostatin for inhibition of the parasite growth. In the absence of PfRKIP, the net concentration of locostatin available for HsRKIP modification may be slightly higher than the corresponding concentrations under normal conditions. This may result in modest increase in parasite sensitivity to locostatin in the *Δpfrkip* parasites. However, the abundant presence of HsRKIP in the RBCs likely mitigates the expected enhancement in the sensitivity of the *Δpfrkip* parasites to locostatin. Therefore, a similar inhibitory profile is observed in the *Δpfrkip* parasites, as in the WT, with the locostatin treatment resulting in a modest, non-significant increase.

Phosphorylation regulates RKIP interactions in multiple contexts. In mammalian systems, S153 phosphorylation drives HsRKIP dimerization and enhances interactions with GRK2 and eIF2α, supporting cellular stress responses^52–54^. PfCDPK1 both phosphorylates and interacts with PfRKIP, modulating their mutual affinity, raising the possibility that PfCDPK1 may function analogously to PKC in regulating PfRKIP in *Plasmodium*.

Although recombinant HsRKIP binds PfCDPK1 with ∼23-fold lower affinity than PfRKIP^10^, the increased import of HsRKIP in *Δpfrkip* parasites may allow functional engagement with PfCDPK1 equivalent to the PfRKIP. Moreover, within the cellular boundaries spatiotemporal changes in the microenvironment surrounding protein assemblies may favour molecular interactions between two proteins that are difficult to recapitulate under *in vitro* conditions. Proximity biotinylation assays with HsRKIP-BioID2 in the *Δpfrkip* background confirmed HsRKIP-PfCDPK1 interactions *in vivo*. Furthermore, in the absence of PfRKIP, HsRKIP forms high a molecular weight complex (100–150 kDa) with PfCDPK1, similar to the PfRKIP-PfCDPK1 complex. Notably, HsRKIP also participates in higher molecular weight assemblies (>150 kDa), suggesting potential roles beyond PfRKIP compensation as supported by import of HsRKIP by the WT parasites in multiple studies including this study^26–28^. HsRKIP may serve as a scaffold facilitating protein-protein interactions, and its phosphorylation by PfCDPK1 further supports functional complementation. The formation of high-molecular weight complexes may restore, in part, the regulatory functions normally provided by PfRKIP, thereby safeguarding the parasite’s invasive capabilities. This observation suggests a compensatory mechanism whereby host RKIP substitutes, at least partially, for the loss of endogenous PfRKIP. It will be interesting to test the growth of *Δpfrkip* parasites in the RBCs devoid of HsRKIP or disrupting PfRKIP in the parasites cultured with RBCs devoid of HsRKIP. However, due to limitations in generating a host RBC devoid of HsRKIP, such experiments may not be feasible^55^.

In summary, our study demonstrates that PfRKIP regulates PfCDPK1 activity through complex formation, which is essential for coordinated phosphorylation events during RBC invasion. Furthermore, this study reveals a novel host-parasite interplay in which the malaria parasite co-opts HsRKIP to maintain essential kinase interactions in the absence of PfRKIP. These findings provide a molecular framework for the development of compounds that enhance PfRKIP-PfCDPK1 interaction, representing a novel strategy to target *P. falciparum* during asexual blood-stage replication. Future studies should focus on delineating the composition of PfRKIP-PfCDPK1 high molecular weight complexes and investigating the regulatory mechanisms by which HsRKIP compensates for PfRKIP loss, including the potential functional consequences of HsRKIP-mediated higher-order assemblies.

## ACKNOWLEDGEMENTS

We thankfully acknowledge Josh Beck for pm2gt-hsp101-bioid2 plasmid, Jacobus Pharmaceutical Company, Inc & Kristin D. Lane for WR99210, Judith L. Green & Anthony A. Holder for the anti-PfCDPK1 antibody. We acknowledge Ashis K. Nandi for allowing the usage of Fluostar Optima (BMG Labtech) plate reader for SYBR Green-I assays. MS received JRF-SRF Fellowship from the Council for Scientific and Industrial Research. We also acknowledge all the lab members who assisted and provided valuable feedback during the course of the study. Inability to include work from other colleagues on the subject is regretted. This work was funded by the Department of Biotechnology (DBT) grant BT/PR28256/MED/29/1313/2018. Some of the work was carried out from another DBT grant BT/PR38411/GET/119/311/2020. Both the grants were awarded to AB. Facilities/laboratories supported by DBT-Builder grant BT/INF/22/SP45382/2022 were utilized for the study. The funding agency has no role in the preparation and decision to publish this work.

## AUTHOR CONTRIBUTIONS

MS generated all the reagents including transgenic parasites; AB, MS designed the experiments; MS performed all the experiments; AB, MS analysed the data and prepared the figures; AB wrote the initial and final manuscript with inputs from MS; All authors reviewed the manuscript.

## DATA AVAILABILITY

All data generated or analysed in this study are included in this manuscript and its Supplementary Information file. The raw data is also available for analysis and reference, if required.

---

**Figure 8.**
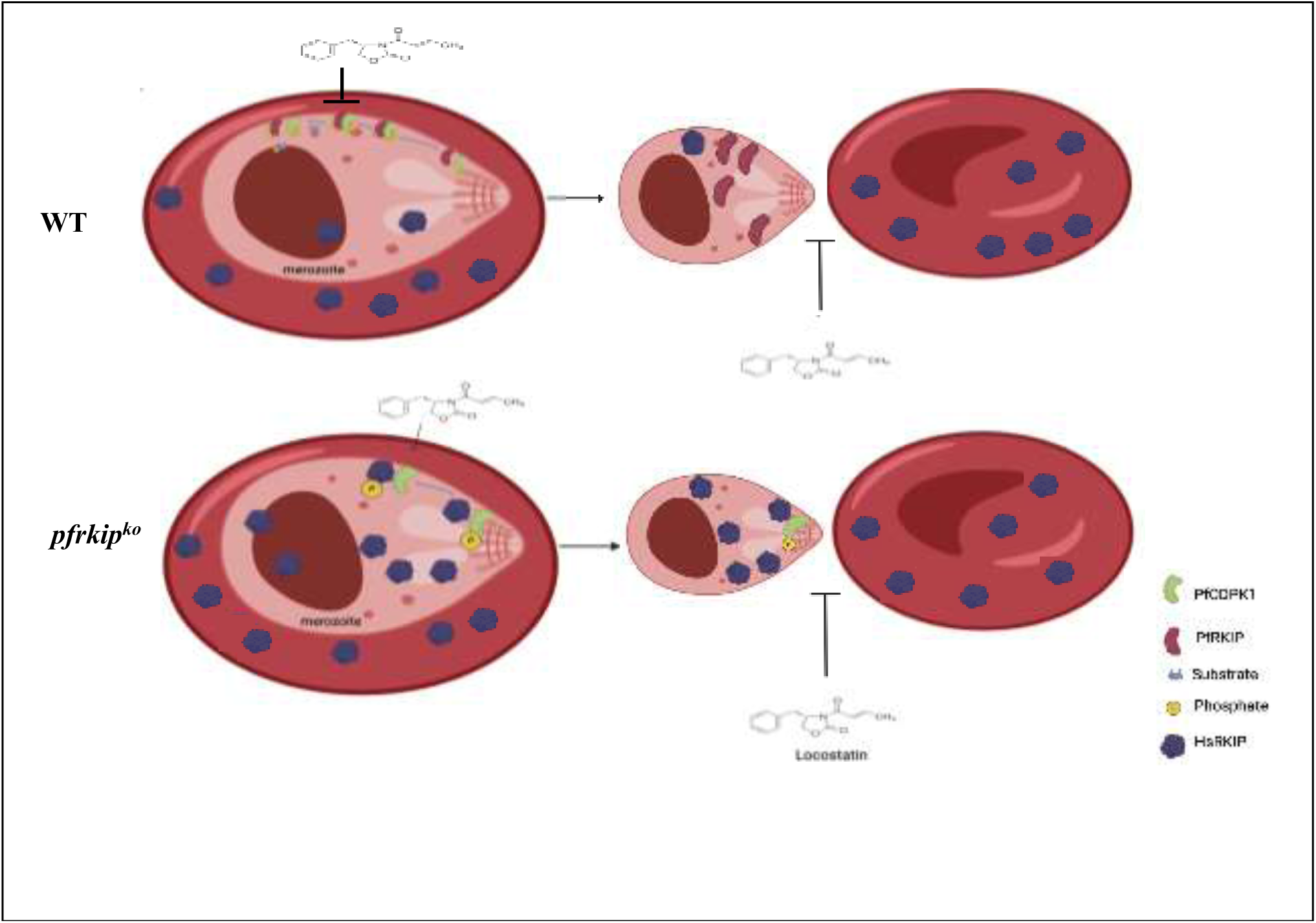
Model illustrating inhibitory activity of locostatin and the adaptive mechanism in *Plasmodium falciparum* following the disruption of *pfrkip*. In wild-type (WT) parasites, PfRKIP modulates the kinase activity of PfCDPK1 through direct interaction^10^. While WT parasites are known to internalize host RKIP (HsRKIP), its functional relevance remains to be elucidated. Locostatin treatment inhibits red blood cell (RBC) invasion in WT parasites, likely by sequestering PfCDPK1 within a stable PfRKIP complex. Abundant host RKIP acts as a sponge reducing the effective concentration of locostatin within the parasite. Notably, genetic knock-out of *pfrkip* (*Δpfrkip*) triggers a compensatory five-fold increase in HsRKIP uptake. In these knockout parasites, HsRKIP substitutes for the absent PfRKIP by interacting with PfCDPK1 to form high-molecular-weight complexes. Consequently, the sensitivity of parasites to locostatin-mediated invasion blockade is maintained, suggesting that PfCDPK1 is perhaps sequestered in complex with locostatin-modified host RKIP as is the case in the WT parasites.

**Figure S1.**
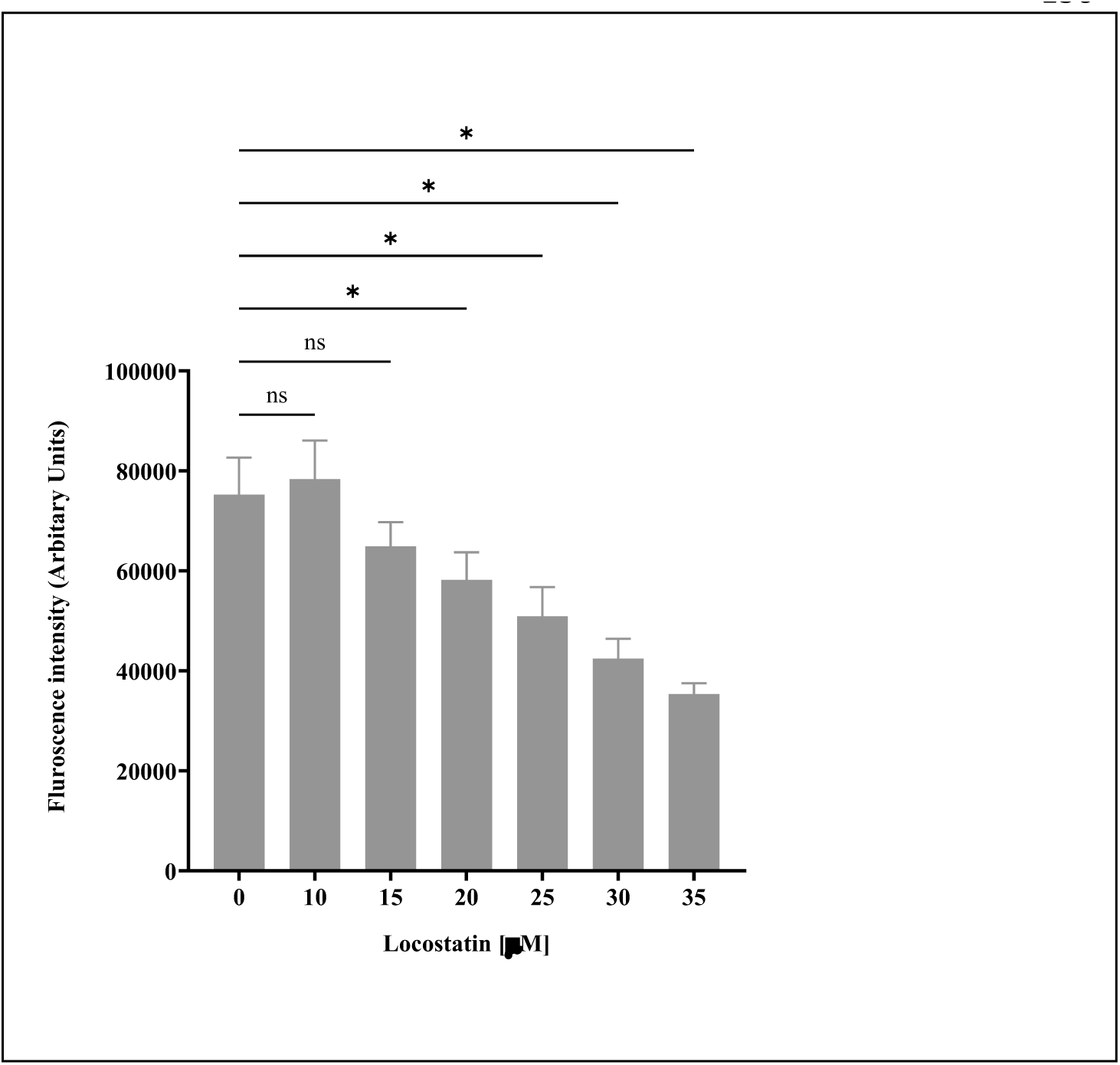
Locostatin inhibits the invasion of red blood cells by *P. falciparum*. To assess the impact of locostatin on the parasite invasion of red blood cells (RBCs), highly synchronized schizonts were treated with increasing concentrations of locostatin. The parasitemia was evaluated after 12–14 h of locostatin treatment through SYBR Green I assay. The parasitemia decreased in a dose-dependent manner with increasing concentrations of locostatin. The fluorescence intensity (arbitrary units) is plotted on the Y-axis against locostatin concentrations (in μM) on the X-axis. The data is obtained from 3 independent biological experiments performed in duplicate (n=3; *p<0.05; ns- not significant, RM one-way ANOVA with Dunnett’s multiple comparisons test). The graph is plotted in GraphPad Prism 10.6.1. The error bars represent the standard error of the mean.

**Figure S2.**
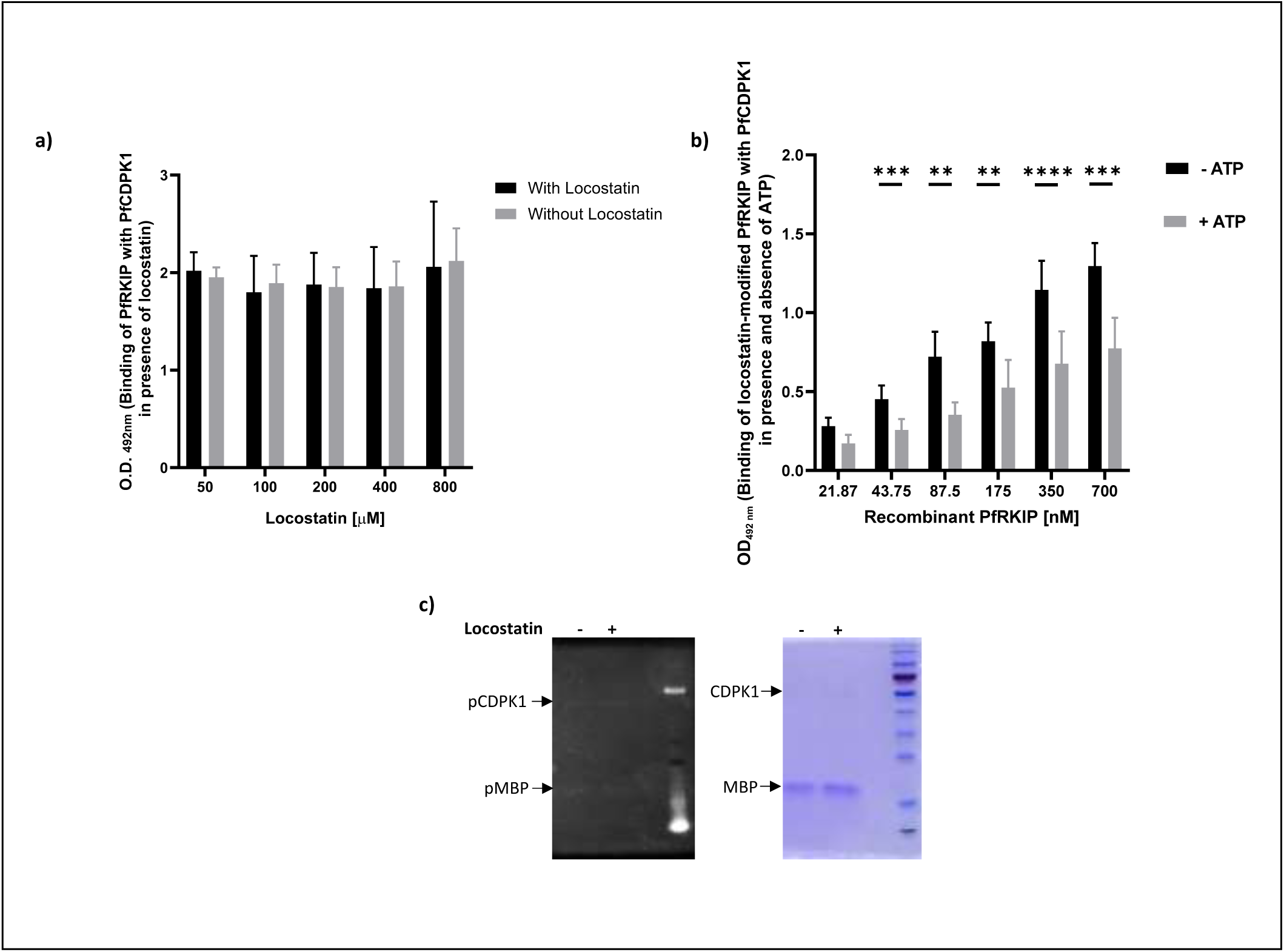
Pre-incubation of PfRKIP with locostatin is essential for enhancing its interaction with PfCDPK1. **a)** Recombinant PfRKIP in the presence of locostatin, without pre-incubation, does not show any change in its interaction with PfCDPK1 compared to without locostatin condition. ELISA was used to coat recombinant PfCDPK1 and incubated with recombinant PfRKIP in the presence or absence of locostatin. The graph shows the concentrations of locostatin (in μM) on the X-axis against OD_492 nm_, representing binding of PfRKIP with PfCDPK1 in presence and absence of locostatin, on the Y-axis. The graph is plotted from data obtained from 2 independent biological experiments performed in duplicate (n=2 biological experiments). GraphPad Prism 10.6.1 is used for plotting the histogram. The error bars represent the standard error of the mean. **b)** In the presence of ATP, the interaction of locostatin-modified PfRKIP with PfCDPK1 is reduced. Recombinant PfRKIP at increasing concentrations was pre-treated with locostatin followed by incubation with PfCDPK1 either in the presence or absence of ATP. The bound PfRKIP was detected with anti-PfRKIP antibodies. Locostatin treated PfRKIP show less interaction with PfCDPK1 in the presence of ATP compared to without ATP condition (n=3; p= 0.0040, paired t-test). The graph shows the concentrations of PfRKIP (in nM) on the X-axis against OD_492 nm_, representing binding of PfRKIP with PfCDPK1 in presence and absence of ATP, on the Y-axis. Error bars represent the standard error of the mean. **c)** Locostatin shows no effect on the kinase activity of PfCDPK1. In vitro kinase assay with recombinant PfCDPK1 and MBP was set up in the presence or absence of locostatin. (left) The reaction products were separated on SDS-PAGE followed by Pro-Q Diamond staining to detect the phosphorylated proteins. The phosphorylation of PfCDPK1 or MBP is the same under the two conditions. (right) The pro-q diamond stained SDS-PAGE gel was counter-stained with Coomassie Brilliant Blue R250.

**Figure S3.**
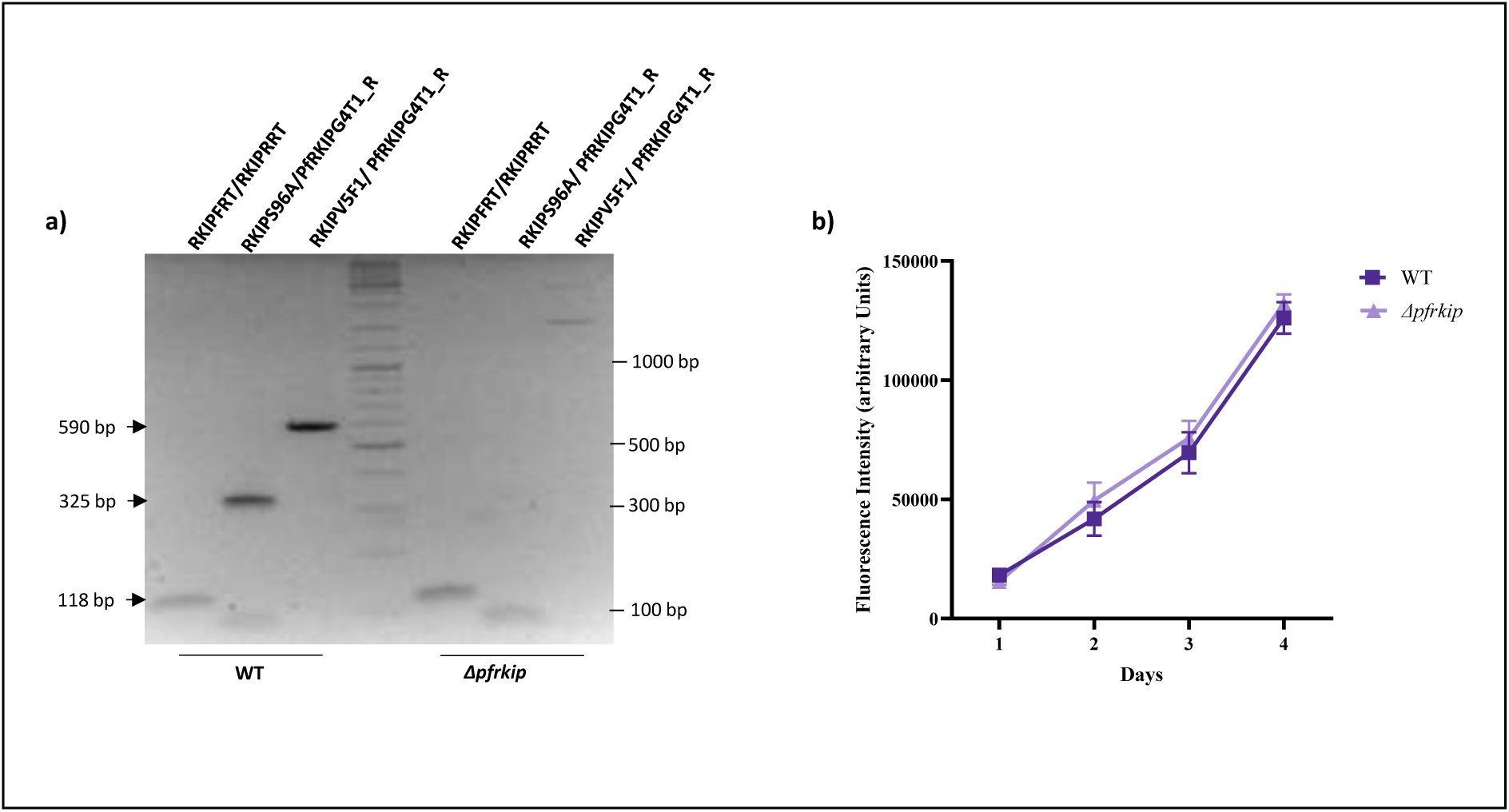
The *pfrkip* knock-out parasites do not express *pfrkip* transcripts and show similar growth as the WT parasites. **a)** Transcripts of *pfrkip* are not expressed in the Δ*pfrkip* parasite. Complementary DNA prepared from the late schizont stage (42–48 h post-invasion) of *pfrkip* and the WT parasites were used as a template to amplify different segments of *pfrkip* using specific primer sets. Remnant segment of *pfrkip* is seen with the RKIPFRT/RKIPRRT primer pair in the Δ*pfrkip* parasites. No amplicons corresponding to the deleted region of pfrkip are seen in the Δ*pfrkip* parasites with the RKIPS96A/PfRKIPG4T1_R and RKIPV5F1/ PfRKIPG4T1_R primer sets. **b)** Asexual growth of the highly synchronized WT and Δ*pfrkip* parasites were compared for two consecutive cycles by SYBR Green I assay. The samples of the WT and Δ*pfrkip* parasites were drawn every 24 h for the period of 4 days and stained with SYBR Green I dye (excitation and emission at 485 nm and 520 nm, respectively). The normalized fluorescence values for the WT and the Δ*pfrkip* parasites are plotted on the Y-axis against the number of days on the X-axis. The growth curve of Δ*pfrkip* parasite closely mirrors the WT parasites, indicating that PfRKIP is not essential for the parasite proliferation under *in vitro* conditions. The error bars represent the standard error of the mean (n=3 independent biological experiments). The graph is plotted using GraphPad Prism version 10.6.1.

**Figure S4.**
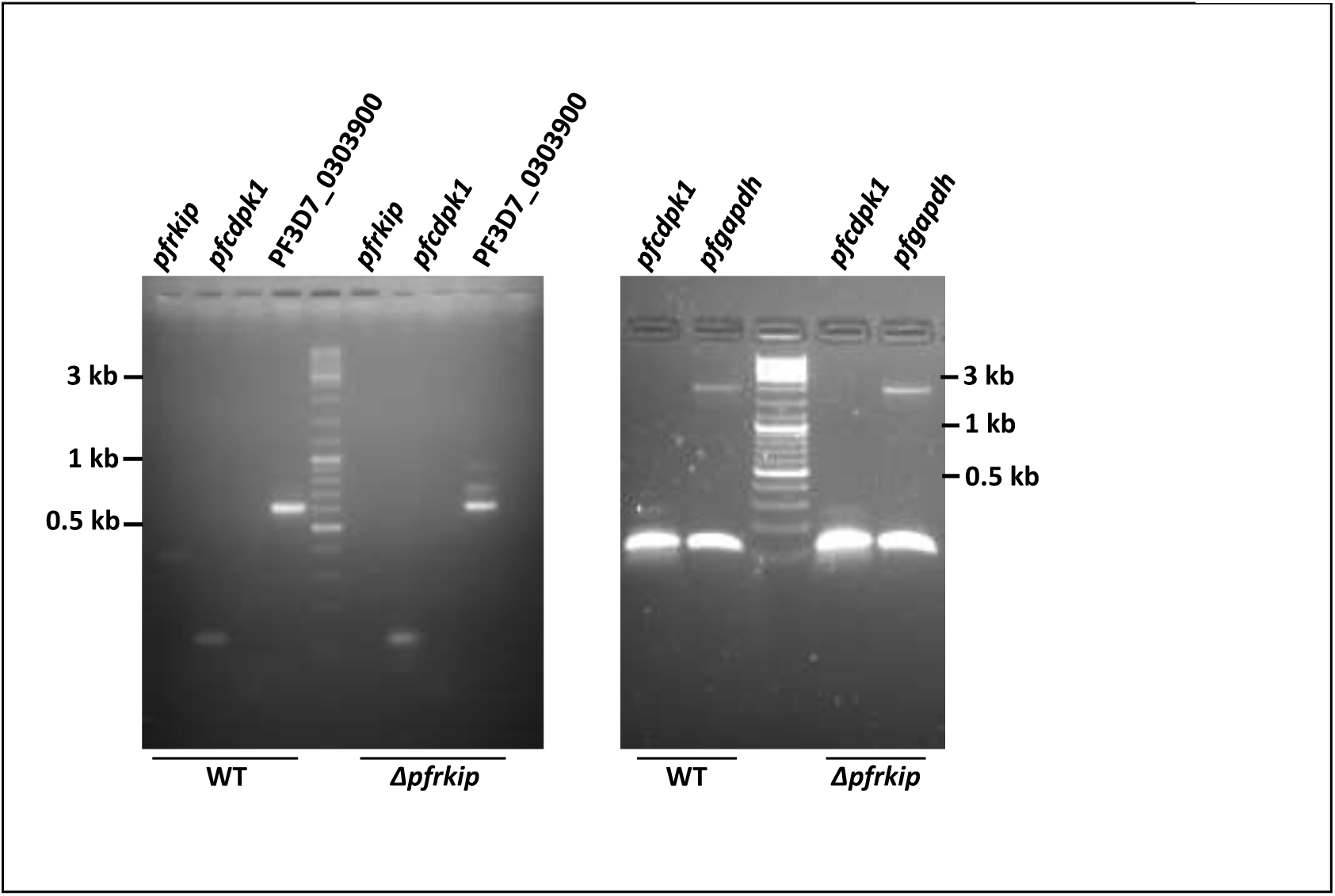
The *pfrkip* knock-out parasites show comparable transcript expression of *pfcdpk1* and PF3D7_0303900 as the WT. The Δ*pfrkip* parasites show similar expression of *pfcdpk1* and PF3D7_0303900 as the WT parasite. To investigate potential compensatory mechanisms in the Δ*pfrkip* parasites in the absence of *pfrkip*, transcript levels of two candidate genes, the PEBP domain-containing protein (Pf3D7_0303900) and PfCDPK1, were measured in both the Δ*pfrkip* and the WT parasites at the late schizont stage (42–48 h post-invasion). (left panel) A house-keeping gene, *pfgapdh*, was used as the internal control. Semi-quantitative PCR analysis show similar expression of *pfcdpk1* and Pf3D7_0303900 transcripts in the Δ*pfrkip* and the WT parasites.

**Figure S5.**
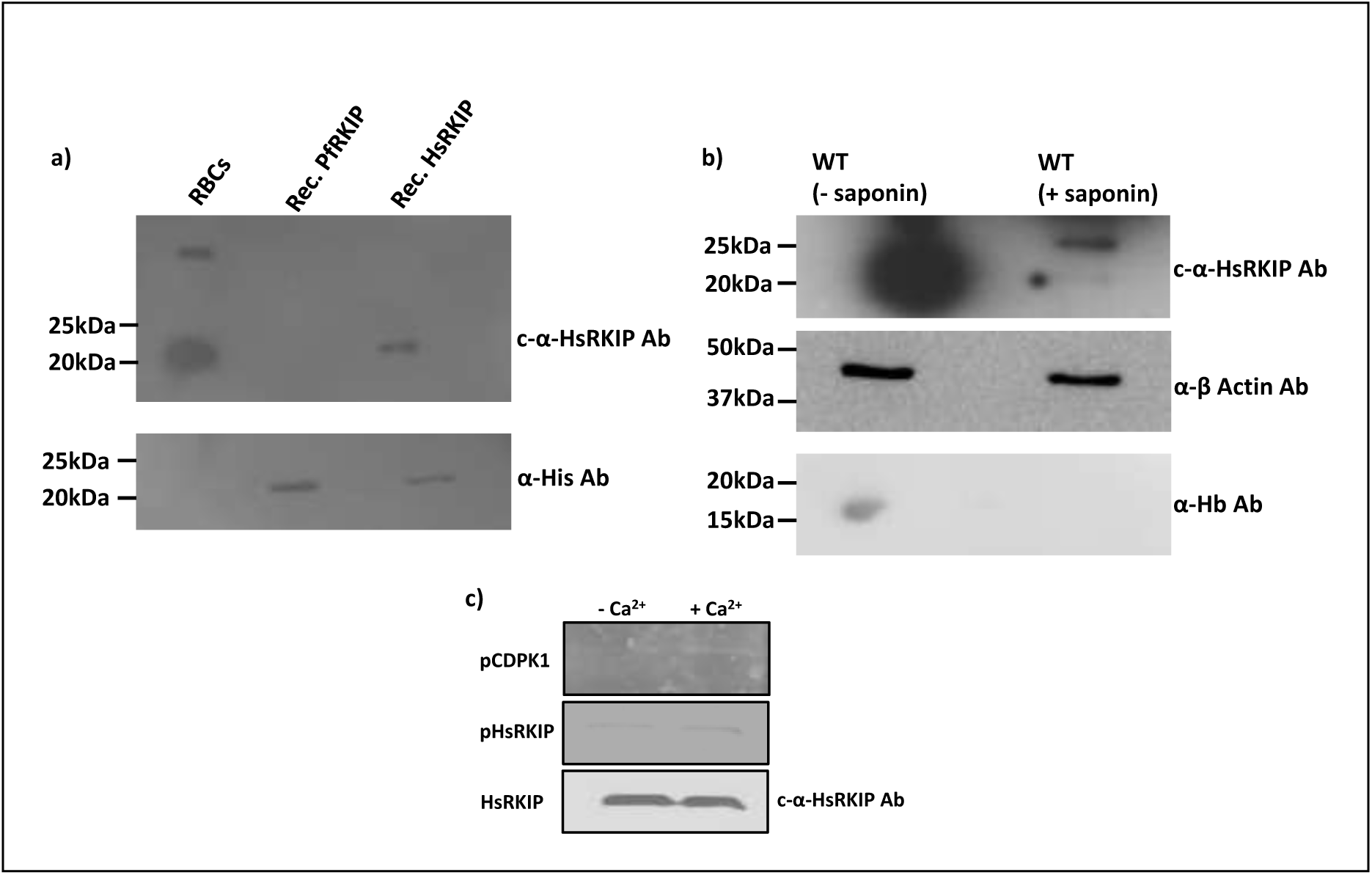
HsRKIP presence in the RBCs and import by the parasite. **a)** Western blot analysis was performed using RBC lysates, recombinant PfRKIP, and recombinant HsRKIP to assess the presence of HsRKIP in RBCs and to validate the specificity of the anti-PEBP1 antibody. The anti-PEBP1 antibody specifically detected HsRKIP in RBC lysates and recombinant HsRKIP at the expected molecular weight of ∼21 kDa but did not recognize recombinant PfRKIP. Loading of recombinant HsRKIP and PfRKIP was confirmed using an anti-His antibody. **b)** Lysates of *P. falciparum*-infected RBCs were fractionated into saponin-lysed fractions (containing parasite compartments only) and non-saponin-treated fractions (containing both parasite and host RBC components). The anti-HsRKIP antibody detected HsRKIP at ∼21 kDa in both the fractions. However, the signal intensity of HsRKIP was substantially less in the saponin-lysed fraction, indicating a lower abundance within the parasite compartments. β-actin was used as a parasite-specific loading control, and anti-Hb antibody confirmed the absence of RBC contamination in the saponin-treated fraction. **c)** Recombinant HsRKIP is phosphorylated by PfCDPK1. In vitro kinase assay with recombinant PfCDPK1 and HsRKIP was performed in the presence and absence of Ca^2+^. The samples were separated on SDS-PAGE followed by Pro-Q Diamond staining to detect phosphorylated products. PfCDPK1 and PfRKIP show calcium dependent increase in phosphorylation. The samples were probed with anti-HsRKIP antibody (c-α-HsRKIP Ab) for loading control.

**Figure S6.**
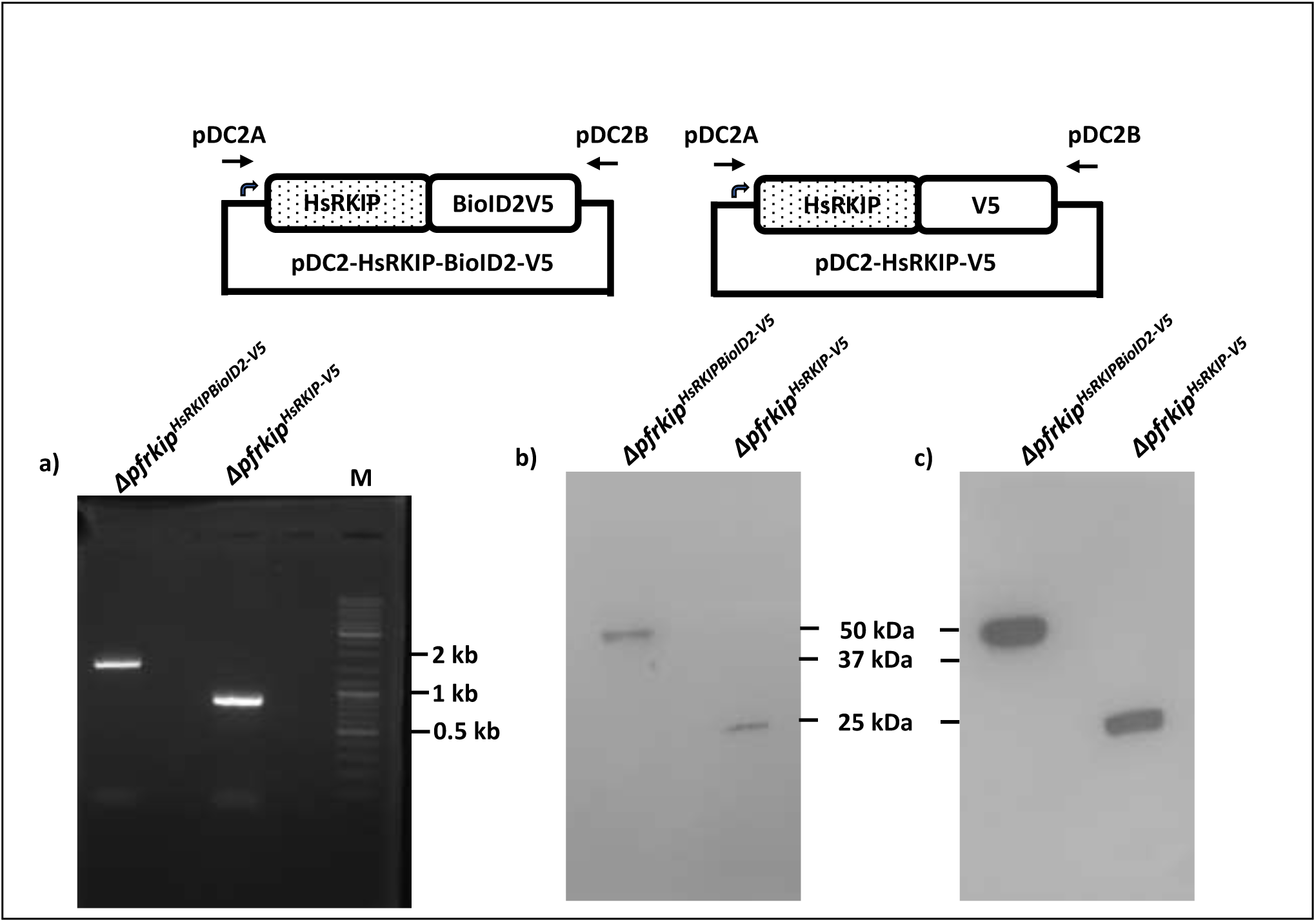
Expression of HsRKIPBioID2-V5 and HsRKIP-V5 in Δ*pfrkip* parasites. **a)** Schematic representation of the plasmid, pDC2, encoding HsRKIPBioID2-V5 and the control plasmid encoding HsRKIP-V5. The HsRKIPBioID2 construct includes a BioID2 tag for biotinylation-based proximity labeling, while the HsRKIPV5 construct serves as a control. **b)** PCR analysis was performed using plasmid-specific primers to validate the presence of pDC2-HsRKIPBioID2-V5 and pDC2-HsRKIP-V5 in the transgenic parasites. The HsRKIPBioID2-V5 expressing parasite, *pfrkip^HsRKIPBioID2-V5^* yielded a specific amplicon of ∼900 bp, corresponding to the BioID2 tag, while the HsRKIPV5-expressing parasite, *pfrkip^HsRKIP-V5^*, produced an amplicon of ∼600 bp. The size shift in *pfrkip^HsRKIPBioID2-V5^* confirms the presence of the BioID2 tag. **c)** The expression of HsRKIPBioID2-V5 and HsRKIP-V5 in *Δpfrkip* parasites was confirmed by Western blotting. Anti-V5 and c-anti-HsRKIP antibodies detected the HsRKIPBioID2-V5 protein at ∼49 kDa and the HsRKIP-V5 protein at ∼23 kDa, respectively, consistent with their predicted sizes. These results confirm the successful expression of both proteins in the *Δpfrkip* background.

**Figure S7.**
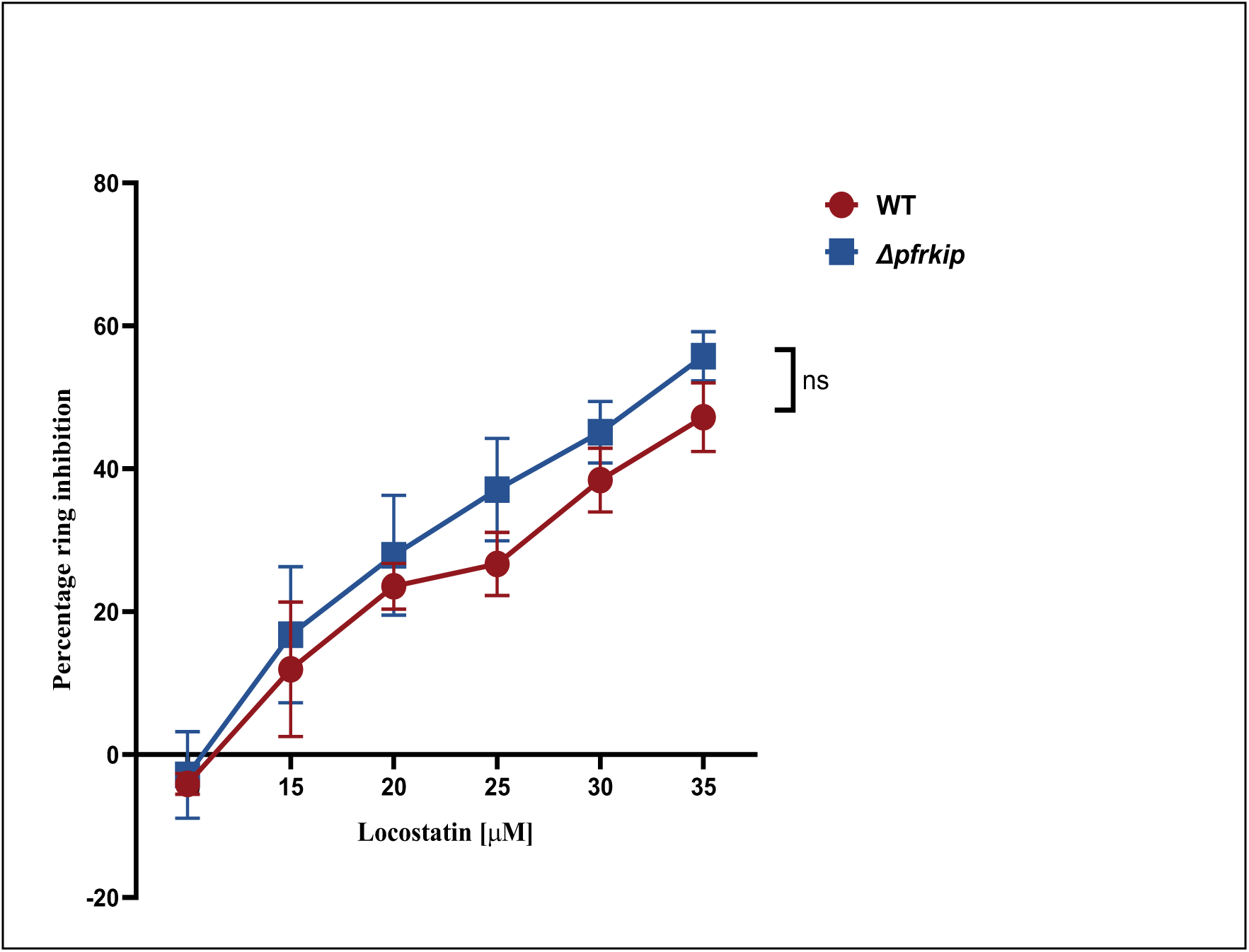
The Δ*pfrkip* parasites show similar sensitivity to locostatin as the WT. Highly synchronized schizonts of Δ*pfrkip* and WT parasites were treated with varying concentrations of locostatin. The percent ring parasitemia was estimated after 12-14 h through SYBR Green I. Locostatin treatment leads to a dose-dependent decrease in the proportion of infected RBCs. The graph shows the percent decrease in ring parasitemia in the WT and the Δ*pfrkip* parasites on the Y-axis at increasing locostatin concentrations [in μM] on the X-axis. Locostatin inhibits both WT and Δ*pfrkip* parasites; however, the Δ*pfrkip* parasites exhibit modestly higher sensitivity to locostatin, as evidenced by a slightly higher inhibition than WT parasites across all tested concentrations. The difference is not statistically significant (n=3, p>0.05, 2way ANOVA, Sidak’s multiple comparison test). Data are expressed as mean ± SEM (n= 3 biological experiments).

**Table 1.**
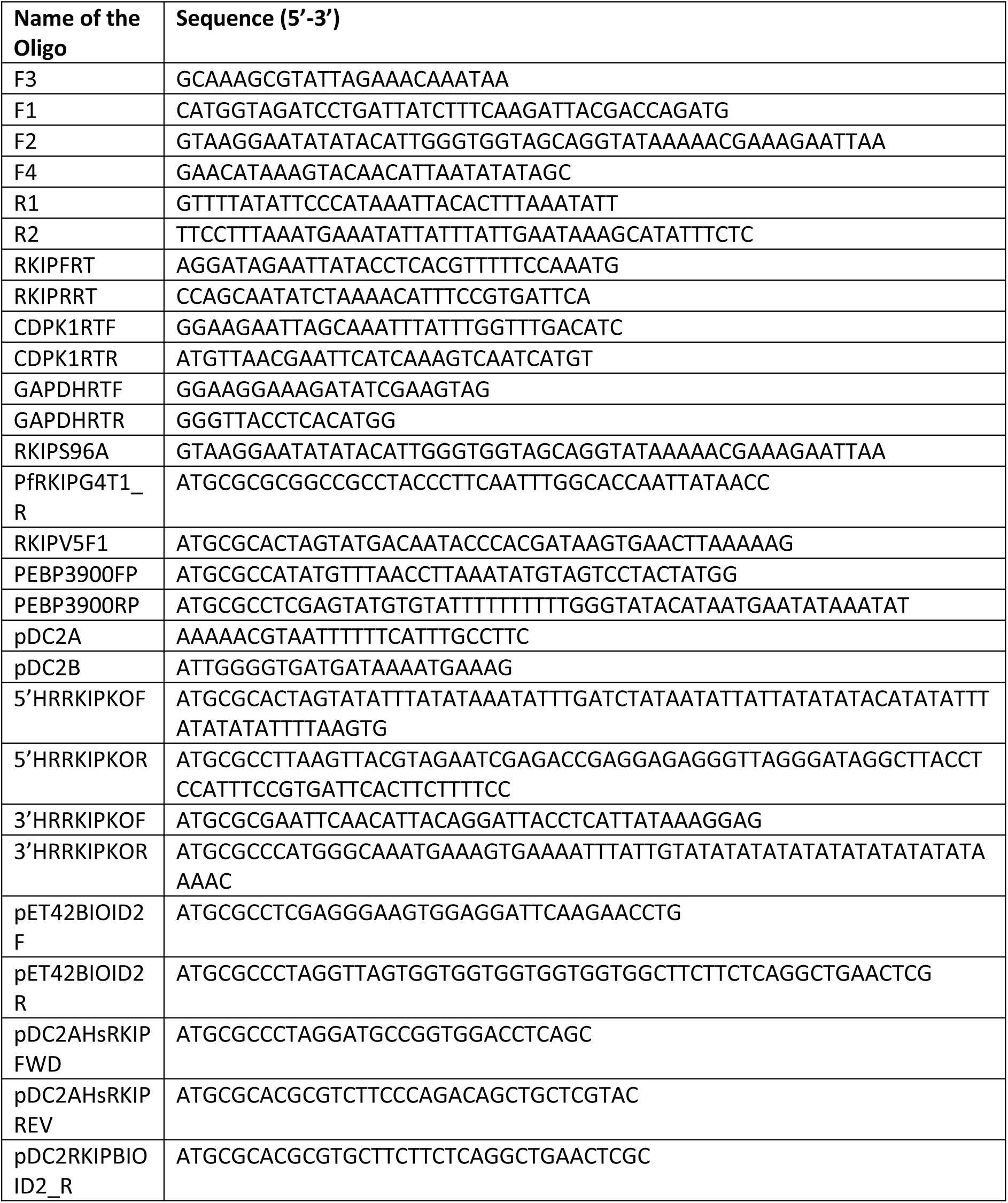
Oligos used in the study.

